# Termination dynamics set RNAPII elongation rate and gate the response to CDK12 inactivation

**DOI:** 10.64898/2026.07.20.737372

**Authors:** Oliver Ozaydin, Lin Liu, Dáire Gannon, Mina Haddawi, Chloe A. N. Gerak, Sarahi Rivera, Laura Corso, Maede Salehi, Mengbo Li, Yining Tian, Tan Nguyen, Jafar Jabbari, Calvin Kraupner-Taylor, Zheng Fan, Jennifer R. Devlin, Traude Beilharz, Gordon K. Smyth, Vihandha Wickramasinghe, John Silke, Ricky W. Johnstone, Benjamin L. Parker, Stephin J. Vervoort

## Abstract

The transcriptional fidelity of RNA polymerase II (RNAPII) is governed by a tight equilibrium between elongation and termination activities, a balance frequently disrupted in human diseases such as cancer. The transcriptional cyclin-dependent kinase 12 (CDK12) maintains RNAPII elongation rate and processivity throughout the gene body. Inactivation of CDK12 disrupts this homeostatic balance and causes elongation stress, slowing RNAPII and triggering premature termination at intronic polyadenylation sites (IPAs). Despite this, the precise executors of intronic premature termination under CDK12-inactivation-induced elongation stress remain poorly understood. Using genome-wide CRISPR screening combined with chemical-genetic approach, we identified a pro-termination mechanism at intronic checkpoints; upon CDK12 inactivation, SCAF4 recruits the cleavage and polyadenylation (CPA) complex through its catalytic endonuclease CPSF3 to execute premature cleavage at IPAs. Disruption of SCAF4-CPA axis prevents early termination, restores full length transcription and confers resistance to CDK12/cyclin K inhibition. Genetic loss of SCAF4 restores the RNAPII elongation rate under CDK12 inhibition, revealing that termination dynamics actively shape the rate of transcription. Supporting this model, we uncovered an anti-termination mechanism driven by KHDRBS1/SAM68, whose depletion promotes proximal termination and sensitises cells to CDK12 targeting. Our findings mechanistically couple elongation and termination activities at intronic checkpoints as joint contributors to both the processivity and elongation rate of RNAPII. This establishes termination dynamics as an active and tractable axis of the cellular response to elongation stress.

## INTRODUCTION

The synthesis of full-length transcripts involves the complex coordination of RNA synthesis with co-transcriptional processing, including splicing and polyadenylation. Maintaining homeostatic balance between RNA Polymerase II (RNAPII)-driven transcription and co-transcriptional processes is critical for cellular homeostasis and the ability to respond to internal and external cues. Subversion of this delicate balance disrupts transcriptional fidelity and can drive human disease including cancer, arising from a failure to create full-length transcripts^1–4^.

The transcriptional fidelity of RNAPII is governed at discrete regulatory checkpoints. These checkpoints, including initiation, pause-release, elongation and termination, are controlled by transcriptional CDKs and their cyclin pairs^1^. At the promoter-proximal pause, the Integrator endonuclease INTS11 terminates a large fraction of RNAPII^5^, opposed by the elongation kinase CDK9/cyclin T and the Integrator-PP2A phosphatase, establishing an active checkpoint that gates entry into productive elongation^6^. During elongation phase, CDK12 and CDK13, in complex with cyclin K, regulate the transcription by directly phosphorylating the RNAPII C-terminal domain (CTD)^7^ and elongation factors such as LEO1^8^. Concordantly, CDK12 inactivation causes widespread elongation stress, characterised by diminished RNAPII processivity, a reduction in the global elongation rate and pronounced premature termination at intronic polyadenylation sites (IPAs)^9–12^. CDK12 loss of function mutations drive genomic instability in cancer^13^, in part through disrupted regulation of DNA damage response genes^9^. Long genes with a high density of intronic polyadenylation motifs are disproportionately sensitive to acute CDK12 inhibition, as this leads to premature transcript cleavage and reduced production of full length mRNA. Since DNA damage response genes are enriched among long, IPAs-rich genes, their preferential truncation under CDK12 inhibition produces a ”BRCAness” phenotype.^9,10^. Crucially, premature termination at IPAs is not unique to CDK12-deficient states and can be detected with slow RNAPII mutants^14,15^, acute PAF1 depletion^16,17^, CDK13 inhibition^11^ and disruption of other elongation factors^17^. The majority of early termination events are detected exclusively under elongation stress, as productively elongating RNAPII in normal conditions bypasses these intronic checkpoints. The fact that the true landscape of intronic termination is only revealed under elongation stress means that the regulatory mechanisms governing this process can only be resolved by studying IPAs directly under conditions of elongation stress, rather than inferred from steady state transcription alone.

Termination at canonical 3’ untranslated regions (UTR) requires the recruitment of the cleavage and polyadenylation (CPA) machinery^18^. This complex is comprised of distinct sub-modules including the cleavage and polyadenylation specificity factor (CPSF), cleavage stimulation factor (CstF), cleavage factor Im (CFIm) and cleavage factor IIm (CFIIm) complexes. Mechanistically, the CPA complex recognises a tripartite poly(A) signal (PAS) containing a canonical AAUAAA hexamer^19–21^, a G/U-rich downstream sequence element bound by the CstF complex^22^ and an upstream UGUA element bound by the CFIm complex^23^. Following assembly, transcript cleavage is executed by CPSF subunits CPSF3 and CPSF100 approximately 10-30 base pairs downstream of the PAS motif^24^, setting the stage for subsequent poly(A) tail addition^25,26^. Fitting with its central role in governing termination, disruption of individual CPA complex members has a widespread impact on transcriptional termination, although the effects can be divergent in both magnitude and directionality^17^.

Elongation and termination activities appear to be reciprocally coupled in cells, with perturbation of either of these processes invariably impacting the other. A key example is telescripting, a process in which the U1 small nuclear ribonucleoprotein (U1 snRNP) complex antagonizes CPA-mediated premature termination by suppressing the utilization of cryptic intronic poly(A) signals, thereby allowing full-length transcript synthesis^27,28^. Central to telescripting is the U1 snRNA, which base-pairs with nascent transcripts at the 5*^′^* splice site to direct the assembly of a U1 CPA factor (U1-CPA complex) thereby sequestering CPA factors from intronic PAS sites.^28^. Consistent with this, genetic abrogation or antisense oligonucleotide (ASO) targeting of the U1 snRNA results in widespread premature cleavage and polyadenylation at IPA sites^27,29^. U1 snRNA targeting however, also invariably impacts the elongation phase of RNAPII with a significantly reduced transcription rate^30^, indicating that both processes are tightly coupled. In line with this notion, combined perturbation of the anti-terminators, SCAF4 and SCAF8, under homeostatic conditions have been demonstrated to cause widespread premature termination at cryptic poly(A) sites, whilst concomitantly reducing the elongation rate of RNAPII^31^. Although the coupling between elongation and termination is well established, its mechanistic basis remains largely unresolved. It is unknown whether termination complexes actively enforce RNAPII deceleration during elongation stress^13,32^. Critically, it is also unknown whether defective elongation can be offset by altered termination dynamics, a possibility that would reframe termination factors as adaptive nodes capable of buffering the elongation deficient states.

Here, we combine genome-wide CRISPR screens with chemical genetics to discover factors that modulate resistance or sensitivity to elongation stress induced by disruption of CDK12. We identify a pro-termination axis, governed by SCAF4-CPA, that operates at intronic checkpoints following CDK12 inactivation. Mechanistically, we showed that upon CDK12 inhibition, the RNAPII CTD interactor SCAF4 recruited CPSF3 to execute premature intronic termination at IPA sites. Loss of this intronic checkpoint recognition, through depletion of SCAF4 or CPSF3, restored full-length transcription despite sustained loss of CDK12 activity. Notably, SCAF4 loss alone was sufficient to restore the RNAPII elongation rate in CDK12-inhibited cells, indicating that intronic checkpoint recognition via this termination factor actively contributes to reduced RNAPII elongation rate. Conversely, loss of KHDRBS1/SAM68, which interact with U1 snRNP to counteract intronic CPA activity, promoted proximal termination and sensitised cells to CDK12 inhibition. Taken together, these discoveries establish that transcriptional fidelity is maintained through the reciprocal integration of elongation and termination factors, and reveal that RNAPII processivity and elongation rate can be set by shifting the balance between these two opposing forces.

## RESULTS

### Genome-wide identification of genetic interactors of CDK12 via chemical-genetics

Canonical termination of transcription at the end of a full transcription cycle is relatively well characterised, as are the molecular mechanisms that govern overall RNAPII elongation. However, the interplay between the two processes remains poorly understood, even though altered elongation nearly inevitably results in diminished processivity and early termination at intronic PAS sites^9,10^. To define the genetic interaction landscape for kinases, we combined, for the first time, genome-wide CRISPR screening with analog-sensitive kinase (AS-kinase) alleles^33^, an approach we term AS-CRISPR-screening. AS-CRISPR-screening negates the need for bespoke small-molecule inhibitors when studying the genetic landscape of a kinase, instead leveraging the rapid and reversible inhibition of catalytic activity via bulky ATP-competitive PP1-analogs such as 3MB-PP1 in AS-kinase engineered cells^34^. To this end, we engineered CDK12-AS and CDK13-AS in adherent HAP1 cells and leukemic suspension cells MV4;11, which exhibit a measurable dependency on CDK12 activity or dual CDK12/13 activity, respectively Fig. S1A Fig. S1B. To validate the HAP1 CDK12^AS/AS^ line for screening applications and characterise the transcriptional impact of CDK12 inhibition, we performed cell proliferation assays, 3*^′^* RNA-sequencing (3’ RNA-seq) and transient-transcriptome sequencing (TT-seq). The results confirmed that acute treatment with 3MB-PP1 led to reduced cellular proliferation Fig. S1C, proximal termination at intronic poly(A) sites Fig. 1A Fig. S1J and diminished processivity of RNAPII exemplified by the *BRIP1* locus Fig. 1A. Consistent with previous reports, CDK12 inhibition also induced the classical BRCAness phenomenon with potent downregulation of BRCA1, FANCD2 and other genes involved in DNA damage regulation^9–11^ Fig. S1H.

**Figure 1.**
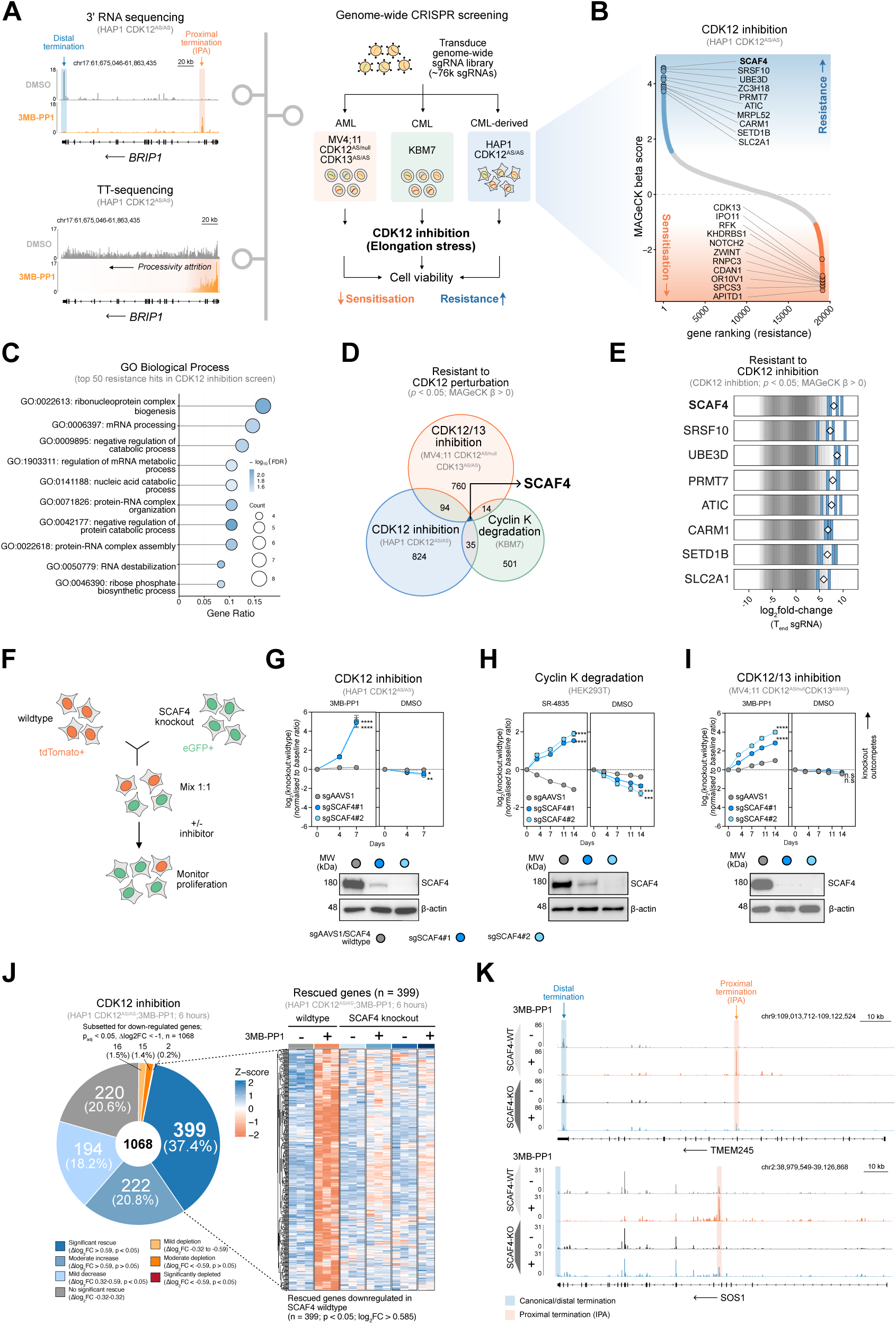
AS-CRISPR-screens reveal that SCAF4 enforces the fitness and transcriptional defects induced by CDK12 inactivation. (A) Left, genome browser view of 3’ RNA-seq and TT-seq signal at *BRIP1* locus on the reverse strand in HAP1 CDK12^AS/AS^ cells treated with 5μM 3MB-PP1 for 6h (3’ RNA-seq) or 30min (TT-seq). Right, schematic of the CRISPR-Cas9 genome-wide screens: CDK12^AS/null^CDK13^AS/AS^MV4;11 cells, KBM7 cells and CDK12^AS/AS^ HAP1 cells were transduced with Brunello library and cultured with DMSO or 3MB-PP1 (150nM MV4;11 and 5μM HAP1) or 20nM SR4835 (KBM7) for 21 days. (B) Genes ranked from highest to lowest MAGeCK *β* scores from genome-wide knockout screens in HAP1 CDK12^AS/AS^ following CDK12 inhibition. The colour of points is indicative of significant resistance (blue) and sensitisation hits (orange). (C) Gene ontology (GO) analysis of the top 50 resistance hits from genome-wide CRISPR screen data in HAP1 CDK12^AS/AS^ cells post-CDK12 inhibition. (D) Venn diagram showing the overlapping resistance hits from genome-wide CRISPR screens with CDK12 perturbation (p < 0.05 and MAGeCK *β* > 0). (E) log_2_fold-change in sgRNA abundance of the top 8 resistance hits following CDK12 inactivation in HAP1 CDK12^AS/AS^ cells. (F) Schematic of the competitive proliferation assays for (G) (H), and (I); SCAF4 wildtype and SCAF4 knockout cells were mixed 1:1 and cultured with vehicle control (DMSO) or CDK12 perturbation. (G) Western blot and competitive proliferation assay for HAP1 CDK12^AS/AS^ cells expressing indicated sgRNAs and treated as indicated. Significance values were calculated through one-way ANOVA and are represented as follows: ns: p > 0.05, *: p *≤* 0.05, **: p *≤* 0.01, ***: p *≤* 0.001, ****: p *≤* 0.0001. Performed in biological and technical triplicate. (H) Western blot and competitive proliferation assay for HEK293T cells expressing indicated sgRNAs and treated as indicated. Significance values were calculated as described in (G). Performed in biological and technical triplicate. (I) Western blot and competitive proliferation assay for MV-4;11 CDK12^AS/null^CDK13^AS/AS^ cells expressing indicated sgRNAs and treated as indicated. Significance values were calculated as described in (G). Performed in biological and technical triplicate. (J) 3’RNA-seq of SCAF4 wildtype and SCAF knockout HAP1 CDK12^AS/AS^cells treated with DMSO or 5μM 3MB-PP1 for 6 hours. Downregulated genes following CDK12 inhibition (n = 1068) are categorised based on their change in the interaction model, with the dark blue segment representing significantly rescued genes (p_adj_ < 0.05, log_2_FC > 0.59). Rescued genes following SCAF4 knockout in are displayed in a subsequent heatmap with cell colour corresponding to z-score log_2_FC per replicate. RNA-seq was performed using three independent technical replicates, except for the SCAF4 knockout + 3MB-PP1 condition, for which one biological replicate failed quality control and was excluded from the analysis. (K) Genome browser view of 3’ RNA-seq signal at *TMEM245* locus and *SOS1* locus on the reverse strand in SCAF4 wildtype and SCAF knockout CDK12^AS/AS^ HAP1 cells treated with 5μM 3MB-PP1 for 6 hours.

Having established a robust model for CDK12 inactivation, we performed genome-wide AS-CRISPR knockout screens in HAP1 CDK12^AS/AS^ cells and MV-4;11 CDK12^AS/null^ CDK13^AS/AS^ treated with 3MB-PP1 and complemented these with CRISPR-screens in KBM7 cells treated with cyclin K degrader SR-4835^35^ Figs. S1B, S1C The resulting data was analysed by MAGeCK and the genes were ranked by MAGeCK *`β`* score to determine genes conferring resistance or sensitisation to CDK12 inhibition or combined CDK12/3 inactivation Figs. 1B, S1D. Functional annotation analysis of top resistance candidates from the HAP1 CDK12^AS/AS^ screen revealed enrichment of factors governing co-transcriptional RNA processing pathways including mRNA processing, ribonucleoprotein complex biogenesis and RNA metabolic pathways Fig. 1C.

To identify the most potent and universal resistance mechanisms, we intersected top resistance hits from the HAP1, MV4;11 and KBM7 screens. SCAF4 was the top hit providing resistance across all three independent screens upon both individual CDK12 inhibition as well as combined CDK12/13 targeting Fig. 1D. Although SCAF4 and SCAF8 are paralogues, we observed no significant enrichment of SCAF8 in any of the screen conditions, nor of the other SCAF family members (SCAF1 and SCAF11) Fig. S1E . This indicates that under the elongation stress induced by inhibition of CDK12 activity, SCAF4 is unique in its resistance mechanism amongst SCAF family members. Consistent with the Gene ontology (GO) analysis in Fig. 1C, the top resistance hits from the HAP1 screen including SCAF4, ZC3H18 and SRSF10 were linked to co-transcriptional RNA processing pathways Fig. 1E, suggesting a direct genetic link between the loss of CDK12 catalytic activity and co-transcriptional processes such as splicing, termination and RNA degradation.

To validate SCAF4 as a resistance candidate, we performed competition assays with tdTomato and eGFP-labelled wild-type and SCAF4 knockout cells, respectively Fig. 1F. Upon CDK12 inhibition in CDK12^AS/AS^ HAP1 cells, SCAF4 knockout cells rapidly out-competed their wild-type counterparts over the course of 7-10 days Fig. 1G. Similar results were obtained in MV4;11 and HEK293 cells with dual CDK12/13 inhibition or CDK12 degradation, respectively. confirming that SCAF4 is required for sensitivity to CDK12 inactivation across different cell lines and targeting modalities Figs. 1H, 1I. Genetic complementation of SCAF4 in the knockout background Figs. S1F, S1G re-sensitised cells to CDK12 inhibition as evidenced by the loss of their competitive advantage over knockout cells in this setting. SCAF4 comprises a well-characterised CTD-interacting domain (CID), which mediates the interaction between SCAF4 and the CTD of RNAPII via Serine2-5 phosphorylated heptad repeats^31^ Fig. S1F. In contrast to the full-length SCAF4 rescue, the CID alone was unable to restore sensitivity to CDK12 inhibition, suggesting that the effect of SCAF4 is not solely driven via competition with other CID-containing proteins Fig. S1G. When ectopically expressed in wild-type cells, the CID domain alone conferred modest levels of resistance, possibly indicating that it interferes with endogenous SCAF4 recruitment Fig. S1G. Re-expression of a mutant lacking the CID domain (SCAF4-ΔCID) drove an intermediate re-sensitisation, a pattern that was mirrored following ectopic expression in wildtype cells Fig. S1G. These data demonstrate that although SCAF4 recruitment to the RNAPII CTD is required for full sensitivity, it is not individually sufficient to drive complete re-sensitisation without the additional contribution of other SCAF4 domains.

To determine the transcriptional consequences of SCAF4 loss following CDK12 inhibition, we next performed 3’ RNA-seq in wild-type and SCAF4 knockout cell lines. Where CDK12 inhibition in SCAF4 wildtype cells led to widespread downregulation of genes, SCAF4 deletion led to a greatly attenuated transcriptional response to CDK12 inhibition. Notably, amongst 1068 downregulated genes upon CDK12 inactivation in HAP1 CDK12^AS/AS^, 399 genes exhibited significant rescue following SCAF4 loss (p_adj_ < 0.05, Δlog2FC ≥ 0.585; 37.4%) Fig. 1J. Experiments in MV4;11 and HEK293T cells with dual CDK12/13 inhibition or CDK12 degradation, respectively, displayed diminished scale and magnitude of the transcriptional response in SCAF4 knockout backgrounds Fig. S1K. Consistent with the rescue in transcriptional response, visualisation of the 3’RNA-seq along CDK12 responsive genes revealed that SCAF4 deletion suppressed CDK12 inhibition-induced IPA events and restored signal in the distal regions around canonical termination sites in the 3’ UTR region of genes Fig. 1K. At steady state, SCAF4 and SCAF8 have been described as functionally redundant anti-termination factors, as proximal polyadenylation occurs only upon the combined loss of both proteins, but not following depletion of either factor alone^31^. Our findings identify a specific requirement for SCAF4 as a pro-terminator executing premature termination at the IPA sites under CDK12 inhibition-induced elongation stress. Rather than contradicting previous work, these findings reconcile SCAF4’s context-dependent functions.

### SCAF4 engages the CPA complex, and reduced CPA engagement or abundance confers CDK12 inhibitor resistance

To determine how SCAF4 can function as an executor of intronic premature termination upon CDK12 inhibition, we next sought to systematically identify the molecular machinery that allows SCAF4 to exert this effect. To this end, we performed comprehensive interactome profiling of SCAF4 using two orthogonal methods, proximity labelling via TurboID-tagged SCAF4 and endogenous SCAF4 co-immunoprecipitation followed by mass spectrometry (IP-MS) Fig. 2A. Across our approaches, SCAF4 was amongst the most enriched proteins, confirming effective bait capture and validating experimental systems before deeper interrogation. Analysis of these datasets revealed robust interactions between SCAF4 and the RNAPII complex, elongation factors such as the PAF1 complex, as well as the core cleavage and polyadenylation (CPA) complex members including CPSF, CstF, CFIm and CFIIm members, amongst others Figs. 2B, 2C Figs. S2A, S2B. These observations are consistent with two distinct domains inherent to SCAF4, the CID and RNA-recognition motif (RRM) domains that mediate interactions with the RNAPII CTD and messenger RNA itself, respectively^31^.

**Figure 2.**
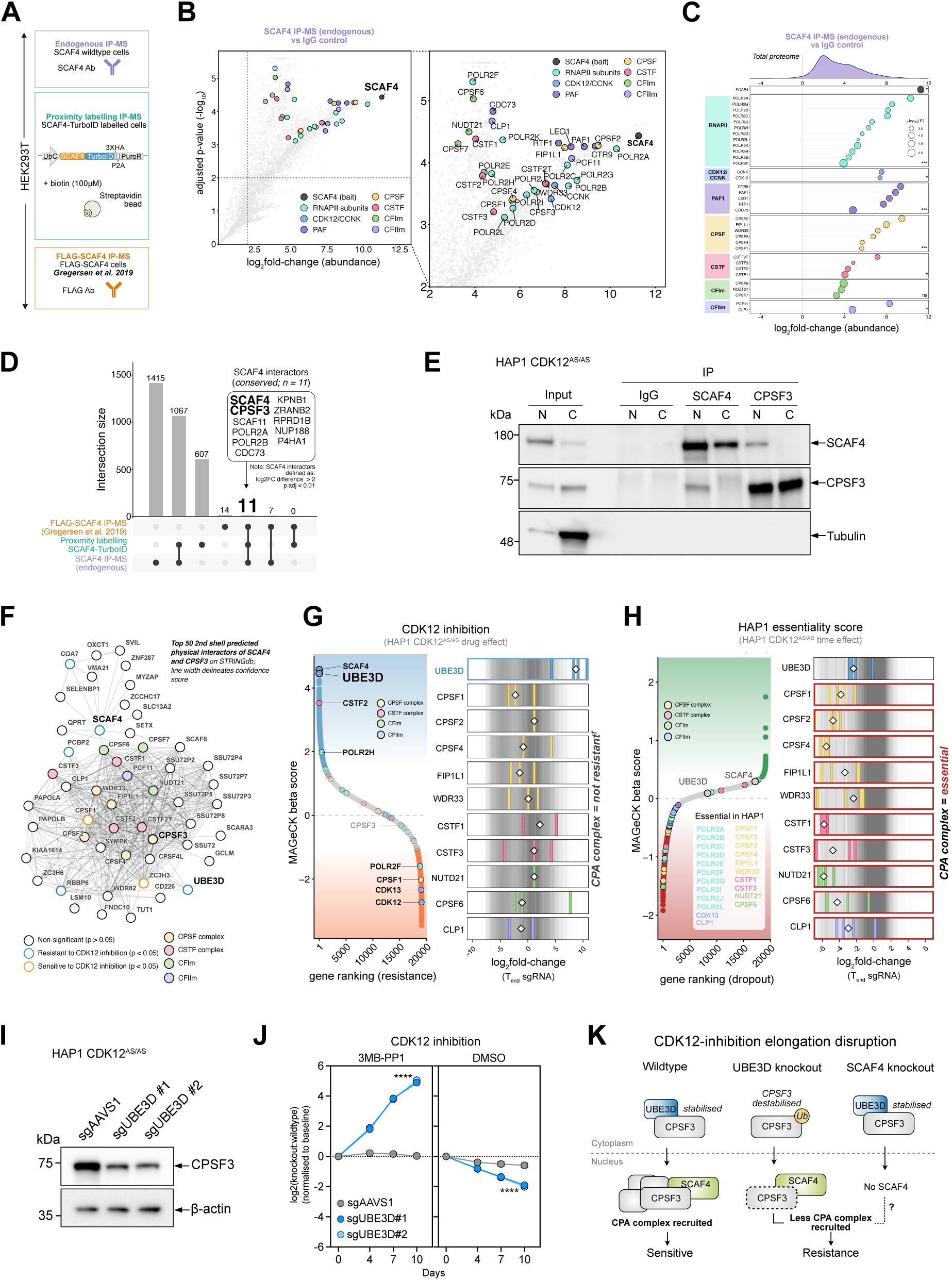
Proteomic profiling uncovers a physical SCAF4-CPSF3 recruitment node linked to UBE3D-mediated resistance. (A) Schematic of the three different MS techniques used to investigate the SCAF4 interactome; Endogenous SCAF4 IP-mass spectrometry (IP-MS), proximity labelling (SCAF4-TurboID), and FLAG-SCAF4-mass spectrometry (IP-MS) from (*Gregersen et al., 2019*)^31^ (B) Volcano plot of quantitative IP-mass spectrometry (IP-MS) of endogenous SCAF4 in HEK293T cells (*n = 4* replicates per condition). (C) Lollipop plot grouped by complex member enrichment in SCAF4 IP-MS data. log_2_fold-change in abundance is indicated on the x-axis, and circle size is indicative of statistical significance. (D) UpSet plot showing the overlap of SCAF4 interactors identified by endogenous SCAF4 IP-MS, SCAF4-TurboID, and FLAG-SCAF4 IP-MS (*Gregersen et al., 2019*)^31^. (E) Reciprocal co-IP Western blot of endogenous SCAF4 and CPSF3 from nuclear (N) and cytoplasmic (C) extracts of HAP1 CDK12^AS/AS^ cells. Immunoprecipitates were probed for SCAF4 and CPSF3. (F) Predicted protein-protein interaction network of the top 50 second-shell interaction between SCAF4 and CPSF3 interactors generated using STRINGdb. Circles are colour-coded by complex with orange outline indicating sensitisation and blue outline indicating resistance to CDK12 inhibition. (G) Left: MAGeCK beta scores ranked by resistance and sensitisation to CDK12 inhibition highlighting SCAF4, UBE3D, RNAPII, CDK12/13 and CPA components. Right: sgRNA abundance for UBE3D and selected CPA factors. Positive log2-fold changes indicate resistance to CDK12 inhibition. (H) Left: MAGeCK beta scores from CDK12^AS/AS^ HAP1 survival screen ranked by gene dropout highlighting RNAPII, CDK13, and CPA components. Right: sgRNA abundance for UBE3D and selected CPA factors at the end time point. (I) Immunoblots for CPSF3 and β-actin from HAP1 CDK12^AS/AS^ cells nucleofected with AAVS1 sgRNA and UBE3D sgRNAs (2 independent sgRNAs). Performed across n = 2 biological replicates. (J) Competitive proliferation assay for HAP1 CDK12^AS/AS^cells expressing indicated sgRNAs and treated as indicated. (K) Proposed model for resistance to CDK12 inhibition in UBE3D and SCAF4 knockout cells.

To distil conserved, high-confidence SCAF4 interactors from our dataset, we cross-referenced our interactors with previously published SCAF4 interactomes derived from FLAG epitope-tagged SCAF4 IP-MS experiments in HEK293T cells^31^. This led to the identification of 11 highly conserved factors, including RNAPII subunits (POLR2A, POLR2B), PAF1 component CDC73, SCAF11 and CPSF complex member CPSF3 Fig. 2D. Notably, CPSF3 (also known as CPSF73), an essential catalytic component of the CPA complex, is responsible for co-transcriptional cleavage of nascent RNA at poly(A) sites^36^. Cleavage of RNA leads to physical, irreversible separation of the nascent transcript from RNAPII, positioning CPSF3 as a plausible, downstream effector of SCAF4-mediated termination at IPA sites. To validate this physical association, we performed reciprocal endogenous co-immunoprecipitation of SCAF4 and CPSF3. Reciprocal immunoblotting confirmed that SCAF4 co-precipitated with CPSF3 in the nuclear fraction and *vice versa*, suggesting that SCAF4 and CPSF3 interact in the nucleus and that SCAF4 may function through the CPA complex Fig. 2E. In line with our findings, factor-specific capture of SCAF4-engaged RNAPII complexes (ELCAP) has independently demonstrated a strong association between SCAF4-bound RNAPII and CPA factors^37^.

To assess if this interaction may be linked to direct CDK12-mediated phosphorylation of CPA members to dynamically modulate CPA recruitment or activation, we intersected high-confidence SCAF4 interactors with phospho-proteomics data following acute CDK12 inhibition Figs. S2C, S2D. CDK12 phospho-substrates were enriched for factors involved in elongation, such as LEO1, as well as splicing and RNA processing factors, but were devoid of obvious direct connections to the core CPA components themselves. This implies that the functional link between CDK12 activity and SCAF4-mediated termination does not rely on direct post-translational modification of the core CPA apparatus, though an indirect regulatory role via upstream elongation or termination factors cannot be completely excluded.

We next reasoned that if SCAF4 exerts its effects via the CPA complex, then genetic inactivation of CPA factors or associated proteins should alter sensitivity to CDK12 inhibition in our CRISPR-screening data. To this end, we first constructed a core CPSF3–SCAF4 interactome using STRING database analysis and expanded it by one additional layer to capture distal interactors Fig. 2F. We then overlaid the CRISPR screen MAGeCK *`β`* scores Fig. 2G and essentiality scores Fig. 2H of these interactors onto this network, thereby highlighting SCAF4-CPA interactors that confer resistance or sensitisation to CDK12 inhibition Fig. 2F. This led to the identification of CPSF3 interactor UBE3D, which, like SCAF4, is non-essential under baseline conditions but emerged as one of the top resistance candidates in CDK12 inactivation CRISPR screens. UBE3D, which was not classified as an E3 ubiquitin ligase in a recent systematic analysis of the human E3 landscape^38^, has been described as a cytoplasmic chaperone of CPSF3 that prevents its proteasomal degradation and facilitates its nuclear import^39^. In addition, we observed resistance via CSTF complex member, CSTF2 and mild sensitisation via CPSF complex member, CPSF1. CPSF1 deletion has recently been demonstrated to cause widespread incorporation of proximal polyadenylation sites in prostate cancer^40^. Beyond this, many of the other CPA complex components did not demonstrate genetic interaction with CDK12 inhibition, including CPSF3 itself. Analysis of the essentiality score revealed that the vast majority of these components were essential under homeostatic conditions Fig. 2H. As core CPA members are indispensable for cell survival, their knockout precludes functional readout in CDK12-inhibited settings.

To validate the UBE3D observation, we generated UBE3D knockout HAP1 CDK12^AS/AS^ cells using single guide RNAs (sgRNAs). Immunobloting revealed that UBE3D deletion led to a pronounced reduction in CPSF3 protein levels, thus driving a cellular state under which CPA complex activity could be considered hypomorphic Fig. 2I. In competitive proliferation assays, UBE3D-deficient cells rapidly outcompeted wild-type cells under CDK12 inhibition, but not under homeostatic conditions Fig. 2J. These data indicate that both impaired recruitment of the CPA following SCAF4 loss and reduced CPA abundance resulting from UBE3D deletion restore cell survival under CDK12-inhibited conditions. This suggests that CPA abundance and SCAF4-mediated CPA recruitment contribute to premature termination and sensitivity to CDK12 inactivation Fig. 2K.

### The SCAF4-CPA axis enforces premature intronic termination to restrict RNAPII processivity under CDK12 inactivation

To study how the disruption of the SCAF4-CPA axis may impact the transcriptional response to CDK12 inactivation we performed 3’RNA-seq in HAP1 CDK12^AS/AS^ cells upon SCAF4, UBE3D or CPSF3 deletion Fig. 3A. Analysis of the resulting datasets revealed over 65% of genes exhibiting significant rescue across one or more conditions Figs. 3B–3C. Consistent with its function as the key catalytic endonuclease in the CPA complex, deletion of CPSF3 had the broadest rescue of gene expression under CDK12 inhibition, closely followed by SCAF4 loss Figs. 3B–3D.

**Figure 3.**
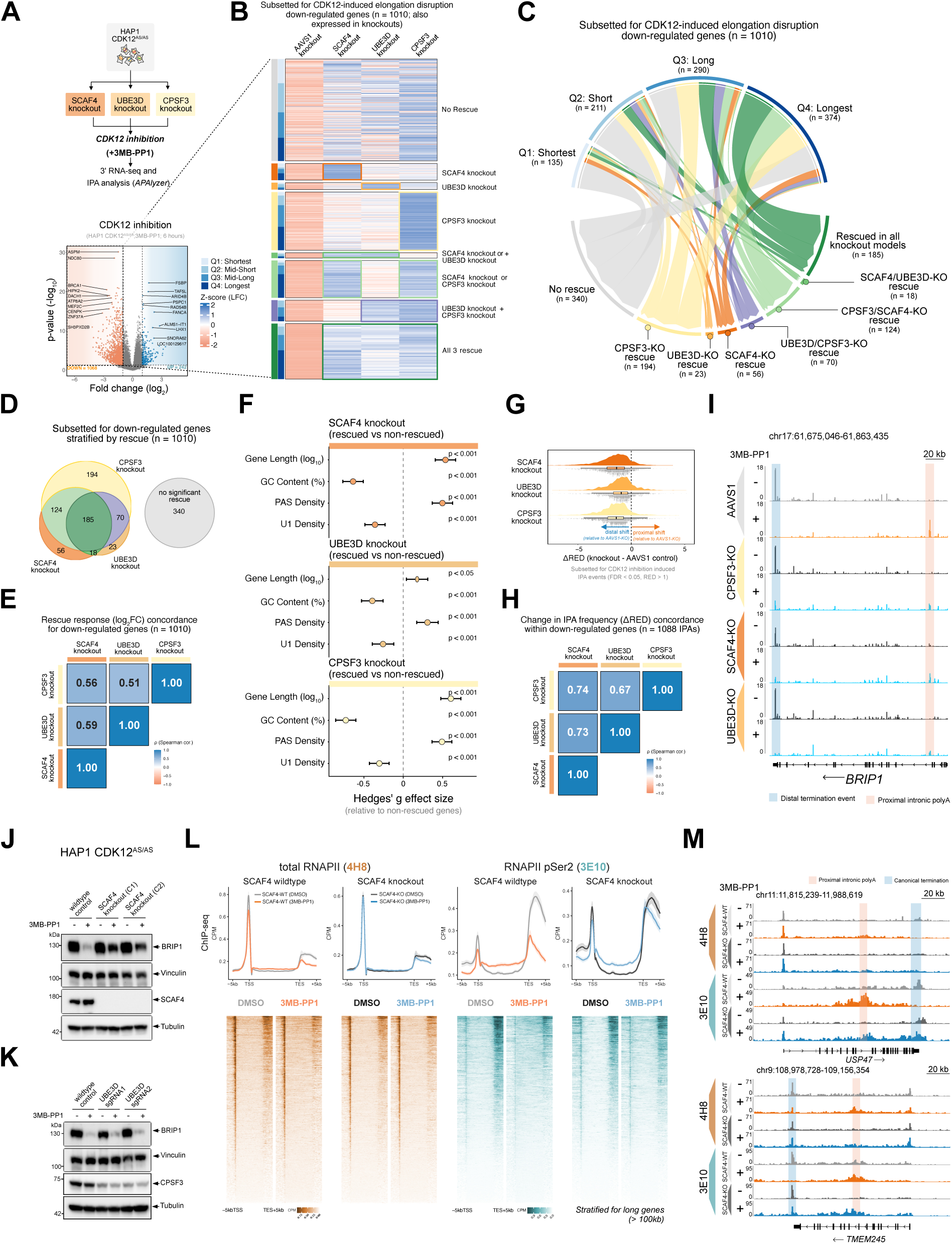
Disruption of a coordinated SCAF4-CPSF3 axis permits RNA polymerase II bypass of intronic checkpoints. (A) Top: Schematic of the 3′ RNA-seq and IPA analysis workflow in HAP1 CDK12^AS/AS^cells following SCAF4, UBE3D, or CPSF3 knockout. Bottom: Volcano plot displaying differentially expressed genes in HAP1 CDK12^AS/AS^cells treated with 5μM 3MB-PP1 for 6h. Significantly up-regulated (*n = 242*) and down-regulated (*n = 1068*) genes are based on a log_2_FC > 1 and adjusted p value (p_adj_ < 0.05). (B) Heatmap of genes downregulated following 6 h of 5 μM 3MB-PP1 treatment in HAP1 CDK12^AS/AS^cells (n = 1,010; genes expressed in all knockout lines). Genes are grouped according to gene length and the knockout genotype(s) in which expression was rescued. Performed across n = 3 technical replicates per condition. (C) Chord diagram of down-regulated genes upon CDK12 inhibition (n = 1,010), grouped by gene length quartile and rescue genotype. (D) Venn diagram showing the overlap of down-regulated genes upon CDK12 inhibition (n = 1,010) rescued by SCAF4, UBE3D, and CPSF3 knockout. (E) Pairwise concordance of rescue responses for down-regulated genes upon CDK12 inhibition (n = 1,010) in SCAF4, UBE3D, and CPSF3 knockout cells. Values indicate Spearman correlation coefficients between rescue response (log_2_FC) in across different genotypes. (F) Hedges’ *g* effect sizes comparing gene length, GC content, PAS density, and U1 density between rescued and non-rescued genes in SCAF4, UBE3D, and CPSF3 knockout cells. Significance values are calculated through two-tailed Wilcoxon rank-sum tests (Mann-Whitney U tests) between rescued and non-rescued gene cohorts. Horizontal error bars represent 95% confidence intervals around the Hedges’ g point estimate. P-values categorised by significance thresholds (p < 0.05, p < 0.01, p < 0.001). (G) Ridge plots showing ΔRED (Relative Expression Difference) values for CDK12 inhibition-induced IPA events (FDR < 0.05, RED > 1) in SCAF4, UBE3D, and CPSF3 knockout cells relative to AAVS1 knockout controls. ΔRED reflects changes in polyadenylation site usage, with negative and positive values indicating distal and proximal shifts, respectively. (H) Pairwise concordance of IPA frequency change (ΔRED) following SCAF4, UBE3D, and CPSF3 knockout. Values indicate Spearman correlation coefficients of ΔRED values in indicated genotypes for 1088 IPA events within down-regulated genes upon CDK12 inhibition. (I) Genome browser view of 3’ RNA-seq signal at *BRIP1* locus on the reverse strand in HAP1 CDK12^AS/AS^ cells treated with 5μM 3MB-PP1 for 6h. Proximal and distal termination sites are highlighted with corresponding scale bar. (J) Immunoblots for BRIP1, Vinculin, SCAF4 and Tubulin from HAP1 CDK12^AS/AS^ cells nucleofected with AAVS1 sgRNA and SCAF4 sgRNAs (2 independent sgRNAs). Performed across n = 2 biological replicates. (K) Immunoblots for BRIP1, Vinculin, CPSF3 and Tubulin from HAP1 CDK12^AS/AS^ cells nucleofected with AAVS1 sgRNA and UBE3D sgRNAs (2 independent sgRNAs). Performed across n = 2 biological replicates. (L) Top: Metagene ChIP-seq profiles for total RNAPII (clone 4H8, orange) and RNAPII CTD pSer2 (clone 3E10, green/blue) across long gene bodies (>100kb) in wildtype versus SCAF4 knockout HAP1 CDK12^AS/AS^ cells. Bottom: Corresponding read-density heatmaps for each condition. (M) Representative ChIP-seq gene-track coverage profiles for total RNAPII (4H8) and RNAPII CTD pSer2 (3E10) at *USP47* (top) and *TMEM245* (bottom) gene loci. Tracks compare HAP1 CDK12^AS/AS^ wildtype and knockout cells following 4 hours treatment with 5*µ*M 3MB-PP1 (+) or DMSO (-) treatment. Proximal and distal termination sites are highlighted with a corresponding scale bar.

This substantial overlap between SCAF4 and CPSF3, both at the gene level and in overall magnitude, further substantiates that they converge on the same axis, transcriptionally and molecularly Figs. 3B–3D. Genes rescued by either of the perturbations were strongly enriched for the longest genes in the genome within the CDK12 sensitive subset, which comprises genes that are longer than average. Intersecting all rescued genes across SCAF4, UBE3D and CPSF3 knockout models yielded a conserved set of 185 genes Fig. 3D enriched for processes such as Notch signalling pathway and DNA-templated transcription amongst others Fig. S3C. Notably, DNA damage response signatures were not universally restored except specific BRCAness genes such as *BRIP1* and *FANCF* Fig. S3D.

We next evaluated the similarity in the rescue response across knockout models in greater detail. Spearman correlation analyses demonstrated a moderate-to-strong positive correlation in their rescue signatures (Spearman’s *`ρ`* = 0.51-0.59) Fig. 3E. This was followed by an examination of the gene-level architectural features inherent to rescued genes compared to non-rescued targets. We observed a conserved architectural signature following stratification of rescued genes across knockouts. Rescued genes were longer, more AT-rich, more PAS-dense, and had a lower density of U1-binding motifs than non-rescued counterparts Fig. 3F.The similarity in gene level responses, and sequence features further suggests a unified mode of action for the SCAF4-CPA axis.

To determine whether the loss of SCAF4, UBE3D or CPSF3 rescues transcriptional response upon CDK12 inactivation through altering premature termination probability at intronic PAS sites, we quantified intronic polyadenylation (IPA) frequency changes using a Δ relative expression difference metric (ΔRED)^41^. A uniform distal shift indicative of a collective suppression of premature intronic cleavage and polyadenylation was observed in SCAF4, UBE3D or CPSF3 knockout cells Fig. 3G Fig. S3L, suggesting that the inactivation of SCAF4-CPA axis was sufficient to prevent early termination at the majority of CDK12 induced IPA sites. Similar to the gene-level analysis, we next determined whether IPA suppression was executed concordantly across knockout lines. Assessment of the correlation in ΔRED values across IPA events sensitive to CDK12 inhibition yields stronger concordance than full-length expression rescue signatures across knockout models, with robust positive correlations (Spearman’s *`ρ`* = 0.67-0.74) Fig. 3H. This concordance is exemplified by gene-track coverage profiles at loci such as *BRIP1* Fig. 3I and *SOS1* Fig. S3I, where signal at proximal intronic polyadenylation sites is reduced in favour of signal at the distal termination site under CDK12 inactivating settings across all knockout models. Although loss of SCAF4 and CPSF3 have been demonstrated to cause read-through of canonical 3’UTRs^31,42^, their role in the regulation of IPA events would be obscured by opposing activities of anti-terminators and pro-elongation factors such as CDK12. These results indicate that termination factors regulate intronic premature termination in a context-dependent manner, with their functional role becoming evident only under elongation stress.

Consistent with the restoration of full-length transcription, BRIP1 protein levels were partially restored in SCAF4 and UBE3D knockout cells Figs. 3J, 3K. Complementing these genetic rescue models, acute pharmacological inhibition of CPSF3 via JTE-607^43^ also phenocopied rescue Figs. S3E–S3G, suppressed premature intronic polyadenylation Figs. S3H, S3J, S3L and restored BRIP1 expression levels, albeit to a lesser extent than SCAF4 deletion Fig. S3K.

Although inactivation of the SCAF4-CPA axis reversed termination and gene expression defects caused by CDK12 inhibition, this could be mediated by direct or indirect effects on transcription. To assess the direct impact of SCAF4 deletion on RNAPII occupancy and phosphorylation, we next performed ChIP-seq for an active pan-phosphorylated form of RNAPII (4H8) and the CTD pSer2 form (3E10). In line with established CDK12 biology, inhibition of its activity resulted in reduced RNAPII occupancy towards distal ends of transcriptional units, with greatly reduced pSer2 and total engaged RNAPII (4H8) levels at canonical termination points. The reduction at distal ends was accompanied by increased signals at promoter proximal sites indicative of non-processive RNAPII and early termination Fig. 3L. In contrast to the pronounced effects in wild-type cells, RNAPII occupancy was largely unaffected by CDK12 inhibition in SCAF4 knockout cells, with only a modest reduction in transcriptionally engaged RNAPII (4H8) and pSer2 occupancy at the distal end of genes and minimal shifts within the gene body. Closer inspection of single gene examples further revealed increased pSer2 signal around intronic polyadenylation sites in wild-type cells upon CDK12 inhibition, a feature closely associated with termination of transcription Fig. 3M. The elevated signal of this mark detected by the 3E10 antibody likely reflects the active dephosphorylation of RNAPII CTD Ser5 following intronic checkpoint recognition. This would subsequently promote pSer2-specific 3E10 antibody binding, which possesses a strict preference for mono-phosphorylated Ser2 heptad repeats^44^. The acquisition of intronic pSer2 signal and reduced occupancy of engaged forms of RNAPII following CDK12 inhibition was greatly attenuated in the SCAF4 knockout background, instead mirroring the wild-type state in DMSO treated cells. Taken together, these observations indicate that loss of the SCAF4-CPA axis members restores full-length transcription, alleviates early termination and leads to increased RNAPII processivity, thus restoring transcriptional fidelity in CDK12 perturbed cells.

### SCAF4 functions as a molecular brake to actively gate RNA polymerase II elongation kinetics

To investigate transcription termination dynamics with greater resolution and sensitivity, we next performed transient transcriptome sequencing (TT-seq) Fig. 4A. Metagene analysis was performed on long genes (> 100kb) to assess the impact of CDK12 inhibition on transcription. The results revealed that CDK12 inhibition in wild-type cells led to an accumulation of signal near the TSS with a sharp attrition in signal throughout the gene body following CDK12 inhibition Fig. 4B. Most prominently, TT-seq signal was almost completely removed from the distal ends of the transcriptional units, as well as in the termination zone directly after. In contrast, this processivity defect induced by CDK12 inhibition was greatly suppressed in the SCAF4 knockout background, as cells maintained robust nascent transcript synthesis throughout the gene body and distal ends Fig. 4B. Even at promoter-proximal sites, SCAF4 knockout cells retained higher TT-seq signal compared to wild-type cells, suggesting that termination dynamics are restored even in the first intron where termination probability differs from the rest of the gene owing to differences in GC content and other sequence elements. In DMSO treated conditions, SCAF4 loss resulted in a modest but elevated occupancy across the gene body, distal ends and the termination zone.

**Figure 4.**
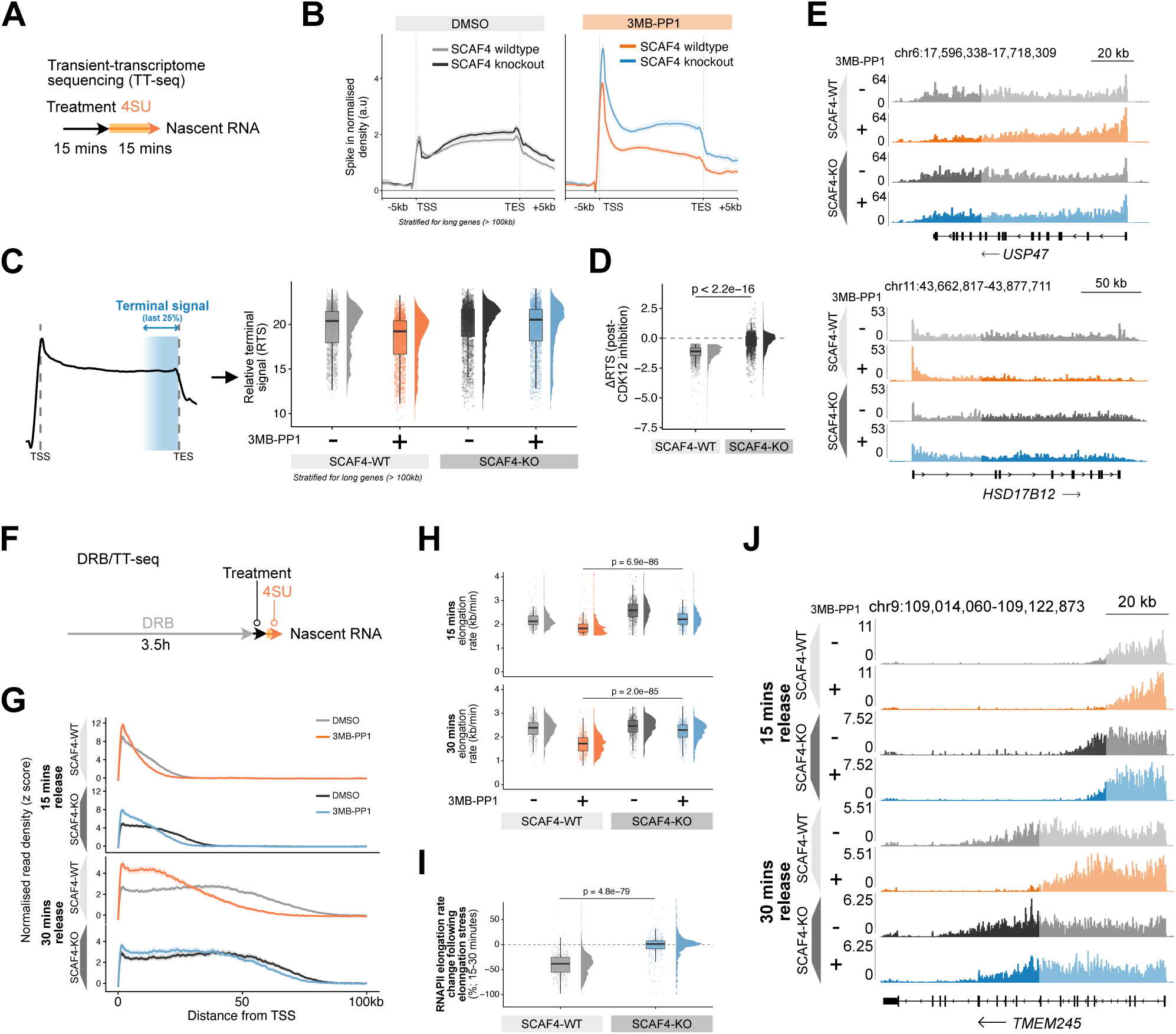
SCAF4 actively gates RNA polymerase II elongation kinetics to enforce intronic checkpoints under CDK12 inactivation. (A) Schematic of the TT-seq workflow in HAP1 CDK12^AS/AS^ cells involving 15 minutes of treatment followed by a 15-minute pulse of 4SU to label nascent RNA. (B) Metagene profiles showing spike-in normalised TT-seq signal intensity across long gene bodies (> 100 kb) in wildtype and SCAF4 knockout cells treated with either DMSO or 10*µ*M 3MB-PP1. The region plotted is scaled relative to the gene body *±* 5 kilobases. (C) Schematic of the relative terminal signal (RTS; last 25% of gene body) metric calculation (left) and raincloud plots showing RTS values across long genes (> 100 kb) in wildtype and SCAF4 knockout cells *±* CDK12 inhibition (right). (D) Raincloud plots of change in RTS to evaluate transcription processivity following CDK12 inhibition in wildtype and SCAF4 knockout backgrounds. p-value was calculated through a two-sample, paired Wilcoxon signed-rank test. (E) Representative genome browser tracks of spike-in normalised TT-seq signal at the *USP47* (top) and *HSD17B12* (bottom) loci across treatment and genotype conditions. (F) Schematic of the DRB/TT-seq protocol used to monitor synchronised transcription wave progression post-DRB release, followed by a 15-minute 4SU pulse. (G) Metagene profiles displaying normalised nascent RNA read density (z-score) as a function of distance from the TSS after 15 or 30 minutes of DRB-release in indicated conditions. (H) Raincloud plots indicating RNAPII elongation rates (kb/min) at 15 min (top) and 30 min (bottom) post-DRB release under DMSO (–) or CDK12-inhibited (+) conditions stratified by genotype. p-values calculated using two-sample, paired Wilcoxon signed-rank test. (I) Raincloud plots representing the percentage change in RNAPII elongation rates between the 15- and 30-minute intervals following CDK12 inhibition per genotype. p-value calculated using a two-sample, paired Wilcoxon signed-rank test. (J) Representative genome browser tracks displaying DRB/TT-seq nascent RNA wavefront progression at the *TMEM245* locus across various conditions and timepoints.

To quantify the TT-seq data, we calculated a relative termination signal (RTS; relative signal in last 25% of the gene body) on a per-gene basis across different conditions. The decreased RTS value in SCAF4 wild-type cells with CDK12 inhibition indicated a severe defect in RNAPII processivity, which was largely rescued in the SCAF4 knockout cells Figs. 4C, 4D. Notably, the TT-seq analysis reflects the effect for all long genes, irrespective of their rescue states at the gene-level, indicating that the restoration of RNAPII processivity is widespread, positioning SCAF4 as a global regulator of intronic premature termination. Indeed, these effects were evident in single gene examples, where there was pronounced rescue of nascent transcript synthesis in the distal termination regions following SCAF4 knockout for *USP47* and *HSD17B12* Fig. 4E.

To determine whether SCAF4-mediated recruitment of the CPA complex or the cleavage activity of CPA complex dictates the restoration of RNAPII processivity, we performed orthogonal TT-seq in cells treated with the CPSF3 catalytic inhibitor JTE-607. Inhibition of CPSF3 partially restored the relative premature termination signal following CDK12 inactivation; albeit the magnitude of processivity rescue was more modest when compared to the SCAF4 knockout cells Figs. S4A–S4C. In contrast, JTE-607 treatment under basal conditions exhibited more pronounced readthrough past the canonical termination site at distal ends of the transcriptional unit, resulting in increased signal in the termination zone Fig. S4D^20,31^. These observations indicate that SCAF4 and its recruitment of the CPA complex is largely selective to intronic checkpoints where its activity is opposed by CDK12. That SCAF4 knockout rescues IPA events more completely than CPSF3 inhibition suggests that physical engagement of the CPA machinery, independent of its catalytic activity, itself restrains RNAPII. In this scenario, SCAF4 loss would remove this engagement entirely, whereas CPSF3 inhibition leaves a recruited but catalytically inert complex in place, which may continue to limit processivity.

To determine whether gene-level architectural features dictated rescue at the nascent level, we stratified long genes (> 100kb with CDK12 inactivation processivity defects (ΔRTS < −0.5) into two cohorts: deemed rescued genes (ΔΔRTS ≥ 0.2; suggests a distal signal improvement) or non-rescued genes (ΔΔRTS ≤ 0.2). This threshold was empirically determined to maintain sufficient cohort sizes for statistical comparison in both SCAF4 knockout and CPSF3 inhibitor JTE-607 backgrounds. In line with 3*^′^*RNA-sequencing observations, rescued genes in both settings were more PAS-dense and more AT-rich than non-rescued counterparts Fig. S4E. We next reasoned that RNAPII processivity rescue may be apparent across intronic polyadenylation sites, specifically evaluating whether SCAF4 deletion relieves nascent signal attrition downstream of CDK12-inactivation-sensitive IPA sites. To quantify this, we calculated an IPA traversal index comparing the nascent RNA signal within the region directly proximal to the IPA (−500bp to 0bp) and a defined downstream window (1000bp to 2000bp). This revealed a significant improvement in downstream nascent transcription traversal in SCAF4 knockout cells relative to wildtype counterparts in CDK12 inhibited settings, demonstrating that SCAF4 loss suppresses termination efficacy at intronic polyadenylation sites Fig. S4F.

To investigate whether SCAF4 deletion enhanced processivity beyond termination dynamics alone, we next assessed its impact on the elongation rate of RNAPII using DRB-TTseq assays. This approach synchronises RNAPII near the transcription start site (TSS) using a reversible elongation inhibitor, 5,6-dichloro-1-*`β`*-D-ribofuranosylbenzimidazole (DRB), followed by releasing cells in DRB-free media for precise temporal tracking of the RNAPII wavefront Fig. 4F. Metagene profiling of nascent read density at long genes (> 100kb) revealed that upon CDK12 inactivation, SCAF4 wildtype cells exhibited a pronounced decrease in RNAPII elongation velocity, as indicated by the diminished distance travelled by the RNAPII wave-front post-release from DRB in 3MB-PP1 treatment cells Fig. 4G. In contrast, SCAF4 deletion largely restored the RNAPII elongation rate under CDK12 inhibition, as indicated by a modest decrease in the distance traversed during the release window, compared to DMSO-treated cells Fig. 4G.

To precisely quantify the elongation rate of RNAPII, we measured the position of the wavefront and then divided the distance traversed on a gene-by-gene level by the time post DRB-release. This revealed that SCAF4 loss effectively rescued wavefront progression defects induced by CDK12 inactivation at both 15 and 30 minute timepoints, with only modest impact on the baseline elongation rate in DMSO treated conditions Figs. 4H, 4I. These observations indicate that in wild-type cells SCAF4 contributes to the termination of transcription and slowdown of RNAPII upon CDK12 inhibition. Under homeostatic conditions where CDK12 is fully active, the presence of SCAF4 does not profoundly impact either of these two processes throughout the elongation phase. Nonetheless, SCAF4 may exert its pro-termination activity at canonical 3’UTR sites as previously demonstrated^31^ and contribute to CPA recruitment and the reduced rate of transcription classically associated with termination, exerted by PNUTS-PP1^45^.

To examine whether SCAF4-mediated restoration of RNAPII elongation rate can be attributed to CPA recruitment or catalytic activity, we subsequently performed analogous DRB-TTseq experiments using the CPSF3 inhibitor JTE-607. Compared to SCAF4 loss, CPSF3 inhibition had a modest to minimal effect on the restoration of RNAPII elongation rate upon CDK12 inactivation, indicating that the effect of SCAF4 depletion on RNAPII elongation extends beyond the mere diminished catalytic activity of the CPA. Figs. S4G–S4I. The kinetic rescue of the RNAPII elongation rate after CDK12 inactivation is illustrated at the *TMEM245* locus following the loss of SCAF4 Fig. 4J and the catalytic inhibition of CPSF3 Fig. S4J. These findings demonstrate that SCAF4 and CPA recruitment actively contribute to the decrease in RNAPII transcription rates under elongation stress and that subversion of this axis can restore productive elongation despite the loss of CDK12 activity. This highlights that termination factors together with canonical elongation factors such as CDK12 set the elongation rate of RNAPII at defined intronic checkpoints.

### KHDRBS1 safeguards selective intronic checkpoints to prevent CDK12 inactivation-induced lethality

Having established the SCAF4-CPA-mediated pro-termination arm at intronic checkpoints following CDK12-inhibition, we next asked whether a counterbalancing anti-termination arm could operate to safeguard transcription elongation. To systematically identify anti-termination factors, we revisited the genome-wide CRISPR knockout screens following CDK12 perturbation Figs. 5A, 5B. We additionally incorporated genome-wide screens in CDK12 and CDK13 analog-sensitive MV4;11 cells, respectively. Analysis of these data sets revealed that across diverse cell types and CDK12 perturbations, KHDRBS1 (also known as SAM68) consistently emerged as a conserved sensitiser across four independent CDK12 inactivation screens. Notably, this genetic interaction was restricted to CDK12 perturbations as KHDRBS1 failed to sensitise to CDK13 inhibition alone, indicating that paralogue asymmetry or non-overlapping functions may drive the genetic interactomes of these two factors Fig. S5A. To gain broader insight into pathways governing sensitivity to CDK12 inhibition, we performed gene ontology (GO) on the top 50 sensitisation hits from both HAP1 CDK12^AS/AS^ and MV-4;11 CDK12^AS/null^ screens Fig. 5C. In either setting, processes related to mRNA splicing emerged as key themes driving sensitivity to CDK12 inactivation. Analogous to the resistance screens, these data suggest a broader interplay between CDK12 inhibition and co-transcriptional machinery also underlies sensitisation. As was evident in the resistance screens, we failed to observe strong enrichment for factors directly involved in transcriptional elongation. KHDRBS1 has been associated with alternative splicing and termination via its physical interaction with splicing machinery^46–48^ such as U1 snRNP subunit U1A^49,50^.

**Figure 5.**
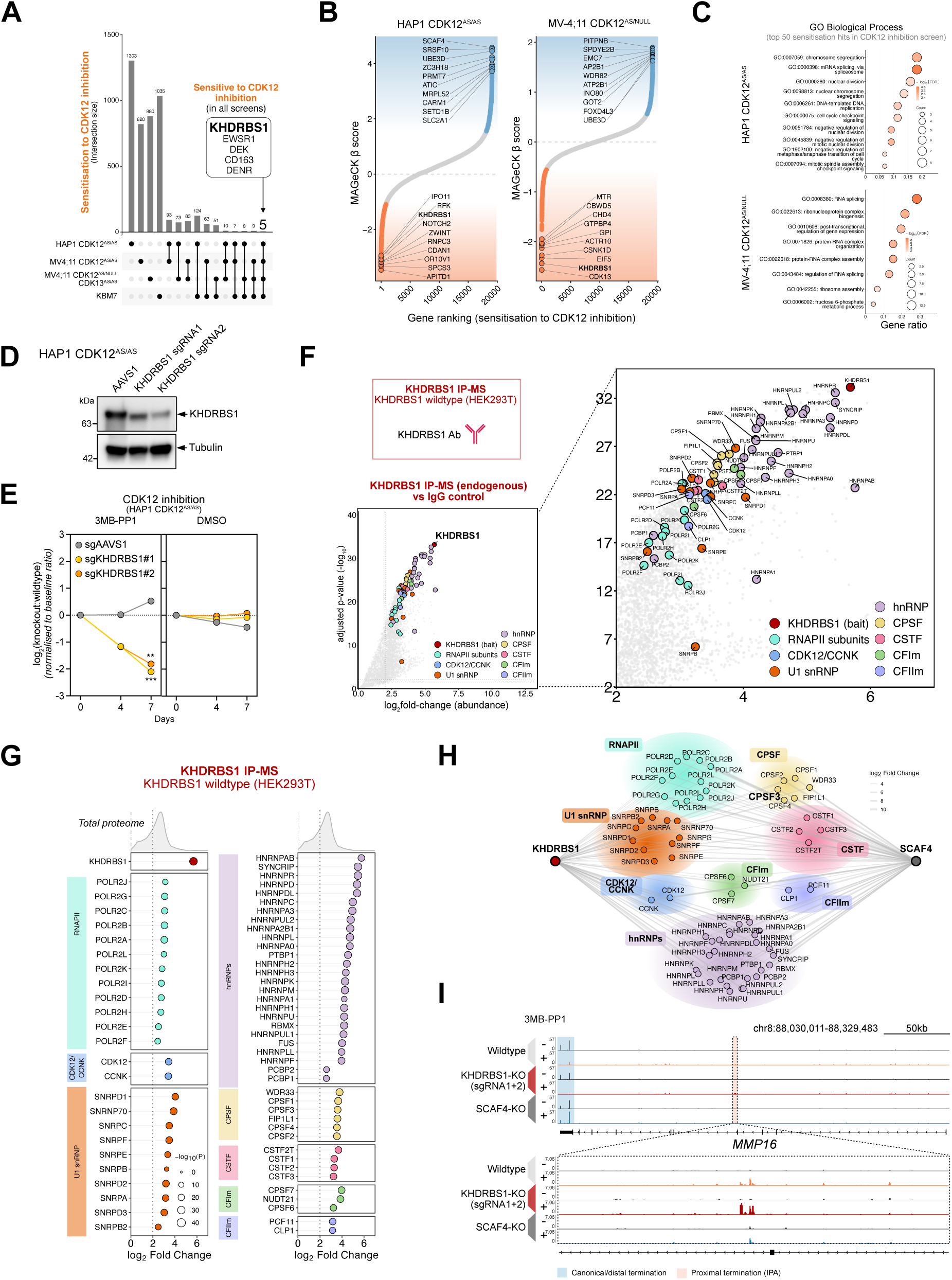
KHDRBS1 acts as an antagonistic rheostat to safeguard processivity at select intronic checkpoints, prohibiting transcriptional collapse and cell death. (A) UpSet plot showing the intersection of sensitisation candidates across genome-wide CRISPR screens, highlighting hits consistent across all screens in the text box. (B) Genes ranked from lowest to highest MAGeCK *β* scores from genome-wide knockout screens in HAP1 CDK12^AS/AS^ (left) and MV-4;11 CDK12^AS/null^ (right) following CDK12 inhibition. Point colour is indicative of significant resistance (blue) and sensitisation hits (orange) (C) Over-representation analysis of the top 50 sensitisation hits from genome-wide CRISPR screen data in HAP1 CDK12^AS/AS^ cells (top) and MV-4;11 CDK12^AS/null^ (bottom) post-CDK12 inhibition.. (D) Western blot validating KHDRBS1 depletion with two independent sgRNAs relative to an AAVS1 sgRNA control, with tubulin as a loading control. Performed across n = 2 biological replicates. (E) Competitive proliferation assays in HAP1 CDK12^AS/AS^ across AAVS1 sgRNA control and two independent KHDRBS1 sgRNA populations following treatment with 5*µ*M 3MB-PP1 (left) or DMSO (right). Significance values were calculated through one-way ANOVA and are represented as follows: ns: p > 0.05, *: p *≤* 0.05, **: p *≤* 0.01, ***: p *≤* 0.001, ****: p *≤* 0.0001. Performed in biological and technical triplicate (n = 3). (F) Volcano plots mapping KHDRBS1 interacting partners identified via endogenous KHDRBS1 immunoprecipitation-mass spectrometry (IP-MS) compared to IgG control, highlighting complexes by colour and corresponding interaction partner names. (G) Lollipop plot grouped by complex member enrichment in KHDRBS1 IP-MS data. log_2_fold-change in abundance is indicated on the x-axis, and circle size is indicative of statistical significance. (H) Protein-protein interaction network diagram constructed from KHDRBS1 and SCAF4 IP-MS data mapping the physical intermediate complexes connected to both KHDRBS1 and SCAF4. (I) Representative genome browser tracks displaying 3*^′^* RNA-sequencing signal at the *MMP16* locus in SCAF4 knockout and KHDRBS1 knockout data. Zoomed-in region highlights further enrichment of an intronic polyadenylation event at *MMP16* locus in the KHDRBS1 knockout population.

We next performed competitive proliferation assays to validate that KHDRBS1 depletion sensitises cells to CDK12 inhibition. KHDRBS1 knockout populations consistently exhibited pronounced proliferative disadvantage upon acute CDK12 inhibition compared to AAVS1 control groups Figs. 5D, 5E. Given that KHDRBS1 is known to cooperate with U1 snRNP to prevent premature transcript termination^49,50^, we reasoned that KHDRBS1 sensitisation may extend to U1 snRNP complex activity. To assess this, we transiently disrupted U1 snRNP function via U1 anti-sense oligonucleotides (U1-ASO) and monitored real-time cell proliferation over 72 hours using an Incucyte imaging platform. While individual U1-ASO treatment or CDK12 inactivation led to moderate proliferative defects, simultaneous disruption of both led to a synergistic arrest of cell proliferation, with confluence flat-lining through the duration of the assay Fig. S5C. These results demonstrate that KHDRBS1 loss or disruption of U1 snRNP alongside CDK12 inactivation results in synergistic collapse of cellular fitness.

To identify the molecular machinery that may be mediating this sensitization phenomenon, we sought to comprehensively map high-confidence KHDRBS1 interactors with immunoprecipitation mass spectrometry (IP-MS). Endogenous KHDRBS1 IP-MS revealed co-purification with the major RNAPII subunits, CDK12/CCNK, establishing its physical proximity to key regulators of transcription elongation. Beyond this, KHDRBS1 co-purified with U1 snRNP complex members, heterogeneous nuclear ribonucleoproteins (hnRNPs) and key CPA complexes including the CPSF, CstF, CFIm and CFIIm complexes Figs. 5F, 5G, supporting previous findings^28,49–51^. Notably, network mapping of KHDRBS1 IP-MS data revealed parallels with the SCAF4 interactome, with extensive convergence on the same core hnRNPs, RNAPII subunits and 3’end processing complexes Fig. 5H. This extensive overlap suggests that, despite opposing responses to CDK12 inhibition, KHDRBS1 and SCAF4 may operate in a similar molecular niche at the intersection of transcription elongation and termination machinery. Rather than acting through isolated mechanisms, these factors may function as opposing rheostats for intronic checkpoints to balance RNAPII processivity outcomes.

To assess whether physical convergence reflects an antagonistic functional relationship at intronic termination checkpoints, we performed 3*^′^* RNA-sequencing following CDK12 inactivation and/or KHDRBS1 depletion. Genes downregulated by KHDRBS1 deletion alone (n = 186) were highly enriched for the KHDRBS1 bipartite motif, with KHDRBS1 itself representing the most depleted transcript in the dataset Figs. S5E, S5G. Differential expression analysis identified a more selective cohort of 239 genes that were synergistically depleted following concomitant KHDRBS1 depletion and CDK12 inactivation Fig. S5F. Mechanistically, IPA events within these genes exhibited a general, synergistic increase in intronic polyadenylation frequency (p = 9.8 × 10^-6^) Fig. S5H. Furthermore, over 50% of synergistically depleted genes at CDK12 inactivation-sensitive loci coincided with genes rescued by SCAF4 loss Fig. S5I. This antagonistic relationship is exemplified at the *MMP16* locus, where under CDK12 inactivation, SCAF4 loss suppresses premature termination in favour of canonical transcript expression, while KHDRBS1 loss further exacerbates premature termination at an already vulnerable IPA site Fig. 5I.

### Bidirectional kinetic coupling between elongation and termination machinery governs intronic checkpoint progression

Our findings suggest a model whereby intronic checkpoint progression is governed by bidirectional kinetic coupling between transcription elongation and termination machinery Fig. 6. The prevailing model of canonical termination positions RNAPII slowdown as an upstream, allosteric determinant of termination fidelity^45^; however, our data presented here extend this paradigm, demonstrating that termination factors themselves can actively shape RNAPII elongation velocity. Under wildtype conditions, basal elongation rates are sufficient to allow RNAPII progression past proximal IPA sites to achieve canonical termination. However, following CDK12 inactivation, a reduced elongation rate triggers premature termination at intronic checkpoints, enabling locus-specific SCAF4 engagement of the cleavage and polyadenylation (CPA) machinery at sensitive genes. While downstream effectors such as CPSF3 drive widespread, non-specific transcript cleavage, SCAF4 functions as a precise intronic regulator of transcriptional termination. As such, SCAF4 deletion rescues CDK12 inactivation-induced premature termination by bypassing intronic checkpoints, restoring full-length transcription and restoring RNAPII elongation velocity. Where SCAF4 depletion alters the recruitment of the CPA complex, diminished expression of its catalytic subunit CPSF3 via UBE3D deletion similarly rescued the termination defect, and full-length transcription albeit to a lesser extend. These data demonstrate that reduced CPA recruitment and levels at these intronic checkpoints dictate the cell’s transcriptional fidelity by controlling both processivity and elongation. Conversely, KHDRBS1 serves as an anti-termination arm of the checkpoint, where it opposes the CPA complex and suppresses intronic polyadenylation at selected genes. As such, KHDRBS1 loss promotes transcriptional collapse and subsequent cell death under CDK12 inactivation. Ultimately, these findings provide the mechanism underlying widespread intronic polyadenylation after CDK12 inactivation, redefining SCAF4 and KHDRBS1 as opposing rheostats that actively gate intronic checkpoint progression. This reframes the notion that termination factors are strictly downstream responders to RNAPII slowdown and recognition of sequence elements, and instead suggests that they may actively shape elongation velocity. Taken together, these findings provide the trans-acting mechanism that underlies gene-level vulnerabilities inherent to CDK12-inhibition-induced elongation stress.

**Figure 6.**
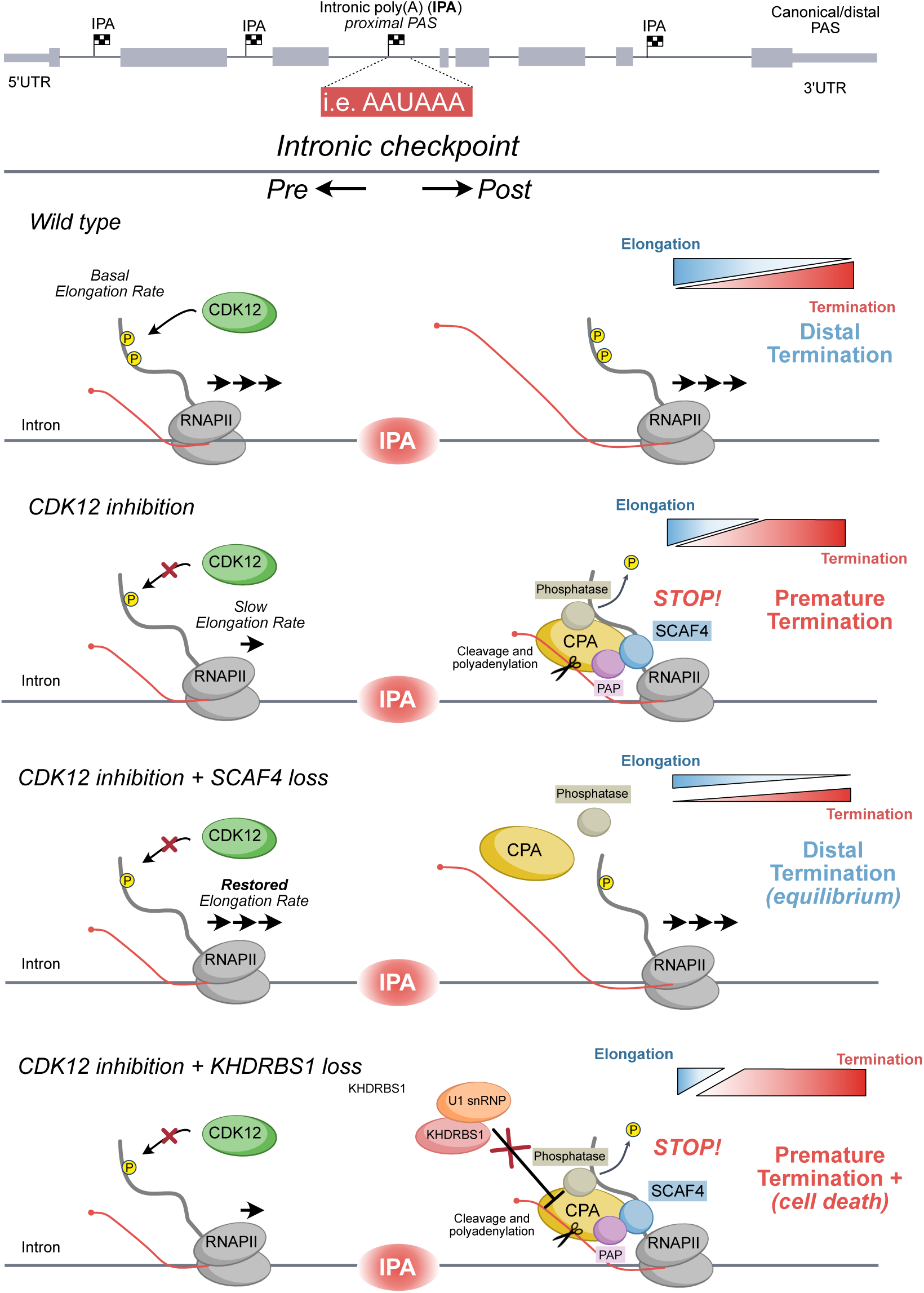
Bidirectional kinetic coupling between elongation and termination factors gates RNAPII termination fidelity at intronic checkpoints. Model describing that intronic checkpoints are governed by bidirectional kinetic coupling between transcription elongation and termination machinery. Under wild-type conditions, basal elongation rates permit RNAPII progression beyond intronic termination checkpoints, resulting in canonical transcript termination. Following CDK12 inactivation, elongation rates are reduced, triggering premature termination at intronic checkpoints in a locus-specific manner, whereby SCAF4 can selectively engage the cleavage and polyadenylation (CPA) machinery. On the other hand, SCAF4 deletion restores RNAPII elongation velocity, bypassing the intronic checkpoint to recover full-length transcription. Conversely, KHDRBS1 serves as a pro-processivity arm of the checkpoint, where it natively opposes the CPA complex and suppresses intronic polyadenylation at select genes. In line with this, KHDRBS1 depletion tips the checkpoint toward premature termination, driving transcriptional collapse and cell death following CDK12 inactivation.

## DISCUSSION

Our findings establish a previously unexplored layer of transcriptional regulation, whereby transcriptional fidelity is maintained by a concerted balance of elongation and termination signals at intronic checkpoints. We establish that opposing CDK12 and SCAF4-CPSF3 activities at intronic polyadenylation sites (IPA) gate RNAPII processivity and set elongation rates, determining the transcriptional fate of long genes. The loss of SCAF4, CPSF3, or the cytoplasmic CPSF3-chaperone, UBE3D restored full-length transcription after CDK12 inactivation, bypassing intronic checkpoints at which RNAPII would otherwise terminate. SCAF4 knockout models demonstrate that loss of termination factors can not only rescue full-length transcription, but actively enhance elongation rates under CDK12-inhibition-induced elongation stress. This work thus identifies SCAF4 as a pro-terminator at intronic checkpoints where it sets the processitivity and elongation rate of RNAPII.

We proposed that under CDK12-inhibition-induced elongation stress, slow RNAPII elongation licenses SCAF4-dependent recognition of nascent RNA at IPA sites, triggering recruitment of the CPA endonuclease CPSF3 to execute intronic premature termination. At canonical 3’UTR sites, the recruitment of the the CPA and associated phosphatases additionally acts to slow RNAPII in the termination zone^45^. We posit similar mechanisms are at play at intronic checkpoints, where the recruitment of the CPA via SCAF4 acts to slow RNAPII creating a direct link between elongation and termination. Our designation of SCAF4 as a pro-terminator does not contradict the loss-of-function phenotypes that originally defined SCAF4 and SCAF8 as anti-terminators under homeostatic states^31^. Indeed, previous work highlighted that SCAF4 depletion alone resulted in modest levels of read-through at canonical 3’UTR sites, in line with a role as a pro-terminator. The pro-termination activity of SCAF4 at intronic sites manifests only under elongation stress, supporting the notion that the functions of pro- and anti-terminators may differ under varying cellular conditions. Our work explains the invariable coupling between elongation and termination and demonstrates that termination factors, together with elongation complexes, set the RNAPII elongation rate.

The relevance of the CDK12-SCAF4-CPA regulatory axis is underscored in human disease characterised by transcriptional fidelity dysregulation. Laboratory models of CDK12 loss broadly exhibit a BRCAness signature involving premature termination and truncation of DNA damage response (DDR) genes, leading to genomic instability and tumourigenesis^9,10,52^. Despite this, patient tumours with bi-allelic CDK12 loss often lack BRCAness signatures observed following acute CDK12 loss and thus are not amenable to PARP inhibitors^13,32^. The lack of a overt BRCAness signature in patients suggest that compensatory mechanisms can offset the loss of CDK12 expression. Possible explanations for such compensation could stem from altered CDK13 activity that has been described as a CDK12 paralog with overlapping functions^11^. Our findings provide an additional mechanism by which cells can adapt to CDK12 loss, namely through the modulation of termination dynamics. It remains to be determined which mechanisms predominate in clinical settings or whether it is a combination of the two. To answer these questions, we believe that future efforts should focus on studying CDK12 function in models that faithfully recapitulate its tumour-suppressive role^53^. Beyond its transcriptional role, the sensitisation mediated by KHDRBS1 deletion to CDK12 inhibition may additionally be related to a shared convergence on DNA-repair mechanisms, as KHDRBS1 has been demonstrated to be involved in PAR-dependent DNA repair which renders KO models sensitive to DNA-damaging agents^54^.

The biological reach of the elongation-termination regulatory axis at intronic checkpoints likely extends beyond CDK12 LOF cancers. The progressive attrition of long gene transcription is a characteristic hallmark of ageing^55–57^. It is plausible that the decline in processivity as we age is governed by a shift in competitive balance between elongation and termination factors. Indeed, SCAF4 is a primary gatekeeper at intronic checkpoints under elongation stress, but it does not restore intronic polyadenylation events universally, potentially suggesting the existence of distinct gene circuits. For instance, SCAF8 and CDK13 are reciprocally predicted to be top co-dependencies according to Cancer Dependency Map data, potentially implying that paralogous factors may govern intronic checkpoints in non-overlapping fractions of the genome, ensuring transcriptional fidelity across gene architectures.

The existence of a CDK12-SCAF4-CPSF3 modulated intronic checkpoint posits parallels between termination in promoter-proximal and intronic regions within gene bodies. CPSF3 and its paralog INTS11 (CPSF3L) are b-CASP-family endonucleases that each terminate RNAPII to gate productive elongation^5,58^. The endonuclease module of the Integrator complex, executes termination at promoter-proximal sites to evict non-productive RNAPII complexes. The Integrator complex also comprises the phosphatase module containing PP2A and INTS6, which dephosphorylate residues within the RNAPII CTD, an activity that works to oppose the pause-release kinase CDK9^5,6^. The intronic checkpoint governed by CDK12 and opposed by a SCAF4-CPSF3 axis thus resembles promoter-proximal control where in both cases the pro-elongation activities of a transcriptional kinase interplay with endonuclease and phosphatase activities to control the fate of RNAPII complexes. The recurrence of regulatory paradigm at promoter-proximal, intronic and canonical 3’ends suggests evolutionary convergence on the same regulatory principles. We propose that sequence dependent checkpoints constitute a recurring architecture that partitions the transcription cycle into discrete, separately controlled phases, and position SCAF4-CPA as the key regulator of intronic termination-elongation checkpoints. Although our screens did not uncover strong enrichment of a phosphatase, their essential nature can obfuscate the identification of a genetic interaction with CDK12 inhibition. At the canonical 3′ end, the CPA complex comprises the PNUTS-PP1 phosphatase which acts to dephosphorylates the RNAPII CTD and in turn decreases the elongation rate ultimately leading to termination and eviction of RNAPII after transcript cleavage^45^. By analogy, the SCAF4-recruited CPA could comprise the same phosphatase module to antagonise CDK12 activity and enforce the intronic checkpoint.

In conclusion, we have established a conceptual framework that leverages bidirectional integration of elongation and termination signals, where opposing CDK12 and SCAF4-CPA activities define plastic intronic checkpoints that modulate transcriptional fidelity under elongation stress. Our work suggested that termination factors are not passive responders to a slowing RNAPII. Rather, they can actively modulate elongation kinetics, and their true function is revealed only when transcription is placed under stress. This provides a plausible mechanistic basis for how biallelic CDK12 loss in patient tumours may restore full-length expression of DDR transcripts, rendering them resistant to PARP inhibition. More broadly, these findings also help define the molecular determination of gene-length-dependent transcriptional fidelity.

### Limitations of this study

In this study, all findings were made in *in vitro* cell line models of acute CDK12 loss. Acute catalytic inhibition may not recapitulate the constitutive loss of CDK12 observed in patients. Adaptive responses to prolonged loss of CDK12 in patients may alter the resistance and sensitisation landscape^13,32^. To discover clinically relevant drug targets, resistance and sensitisation should be studied in patient derived models or mouse models that faithfully recapitulate human disease.

## MATERIALS AND METHODS

### Resource Tables

**Table 1.**
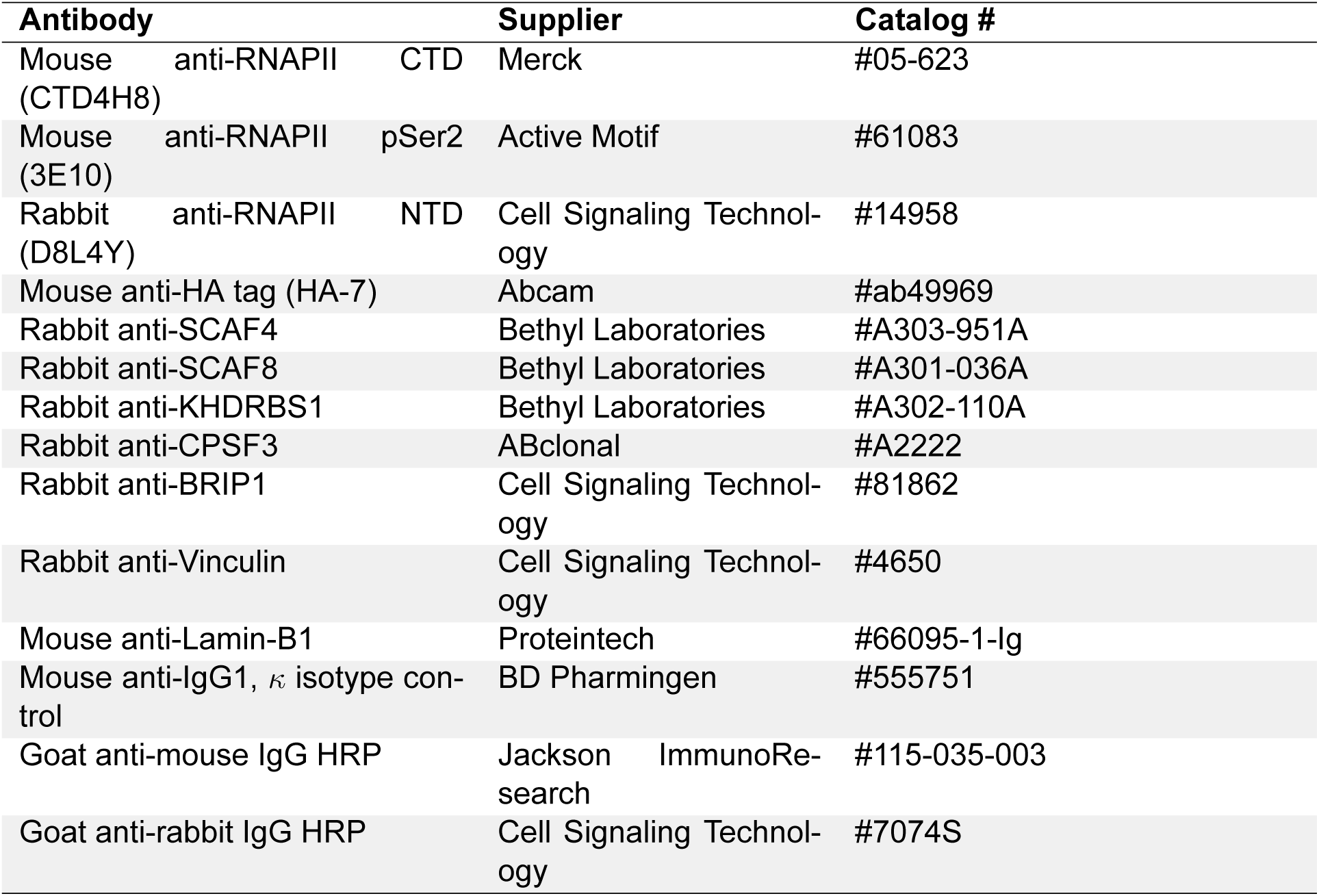
Antibodies used.

**Table 2.** Chemicals, peptides, and recombinant proteins.

| Reagent | Supplier | Catalog # |
| --- | --- | --- |
| 3-MB-PP1 | MedChemExpress | #HY-102069 |
| SR-4835 | MedChemExpress | #HY-130250 |
| JTE-607 | Cayman Chemical | #36942 |
| 5,6-Dichloro-1-β-D-ribofuranosylbenzimidazole (DRB) | Sigma-Aldrich | #D1916 |
| 4-thiouridine (4sU) | Sigma-Aldrich | #T4509 |
| Doxycycline hyclate | Thermo Scientific | #J60579.22 |
| DMSO (suitable for hybridoma) | Sigma-Aldrich | #D2650 |
| UltraPure DNase/RNase free water | Thermo Scientific | #10977015 |
| Penicillin/Streptomycin | Thermo Fisher Scientific | #15140122 |

Table 2 continued from previous page
| Reagent | Supplier | Catalog # |
| --- | --- | --- |
| Foetal bovine serum (Australian origin) | Bovogen Biologicals | #SFBS-AU |
| GlutaMAX™ Supplement | Thermo Fisher Scientific | #35050061 |
| Schneider's Drosophila Medium | Thermo Fisher Scientific | #21720024 |
| MEM Non-Essential Amino Acids Solution | Thermo Fisher Scientific | #11140050 |
| TrypLE™ Express Enzyme (1X) | Thermo Fisher Scientific | #12604021 |
| Lipofectamine 3000 | Invitrogen | #L3000001 |
| Lipofectamine 2000 | Invitrogen | #11668027 |
| Poly(ethyleneimine) solution | Sigma-Aldrich | #03880-500ML |
| Sequa-brene (polybrene) | Sigma-Aldrich | #S2667 |
| Alt-R SpCas9 Nuclease V3 | Integrated DNA Technologies | #1074182 |
| Q5 Ultra II High-Fidelity DNA Polymerase | New England Biolabs | #M0544S |
| XbaI restriction enzyme | New England Biolabs | #R0145S |
| BamHI-HF restriction enzyme | New England Biolabs | #R3136S |
| BsmBI-v2 restriction enzyme | New England Biolabs | #R0739S |
| TRIzol Reagent | Thermo Fisher Scientific | #15596026 |
| Phenol:Chloroform:Isoamyl alcohol (25:24:1) | Thermo Fisher Scientific | #15593031 |
| MTSEA biotin-XX linker | Biotium | #BT90066 |
| HRP-conjugated streptavidin | Thermo Fisher Scientific | #N100 |
| ECL detection reagent | Thermo Fisher Scientific | #34580 |
| μMACS Streptavidin MicroBeads | Miltenyi Biotec | #130-074-101 |
| Protein A/G magnetic beads | Thermo Scientific | #88848 |

**Table 3.** Critical commercial assays.

| Assay | Supplier | Catalogue Number |
| --- | --- | --- |
| *Continued on next page* |  |  |
| μMACS Streptavidin Kit | Miltenyi Biotec | Cat# 130-074-101 |
| Qubit RNA HS Assay Kit | Thermo Fisher | Cat# Q32852 |
| Qubit RNA BR Assay Kit | Thermo Fisher | Cat# Q10210 |
| Qubit DNA HS Assay Kit | Thermo Fisher | Cat# Q32854 |

Table 3 continued from previous page
| Assay | Supplier | Catalogue Number |
| --- | --- | --- |
| Qubit DNA BR Assay Kit | Thermo Fisher | Cat# Q32853 |
| CellTiter-Glo Luminescent Cell Viability Assay | Promega | Cat# G7570 |
| QuantSeq 3' mRNA-Seq V2 Library Prep Kit | Lexogen | Cat# 191.96 |
| Direct-zol™ RNA Purification Miniprep Kit | Zymo Research | Cat# R2052 |
| NEBNext Ultra II Directional RNA Library Prep Kit for Illumina | New England Biolabs | Cat# E7760L) |
| S-Trap™ Universal Sample Prep Micro Kit | ProtiFI | Cat # K002-MICRO-0010K |

**Table 4.** Deposited data.

| Data type | Source | Identifier/Link |
| --- | --- | --- |
| 3' RNA-seq | This paper | GEO: *Will be provided* |
| TT-seq | This paper | GEO: *Will be provided* |
| DRB-TT-seq | This paper | GEO: *Will be provided* |
| Unprocessed and uncompressed imaging data | This paper | Mendeley: *Will be provided* |
| Proteomics data | This paper | PRIDE: *Will be provided* |

**Table 5.**
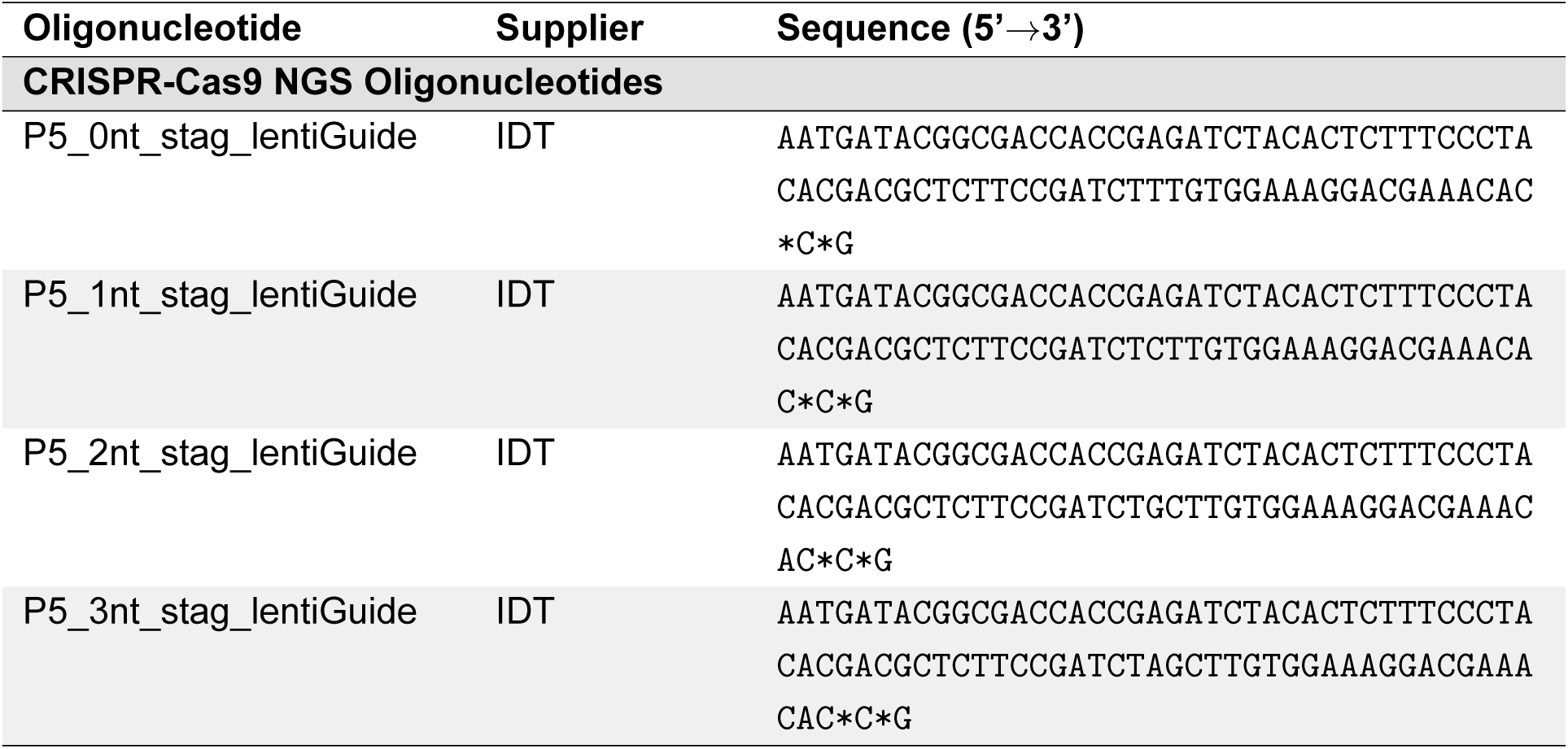
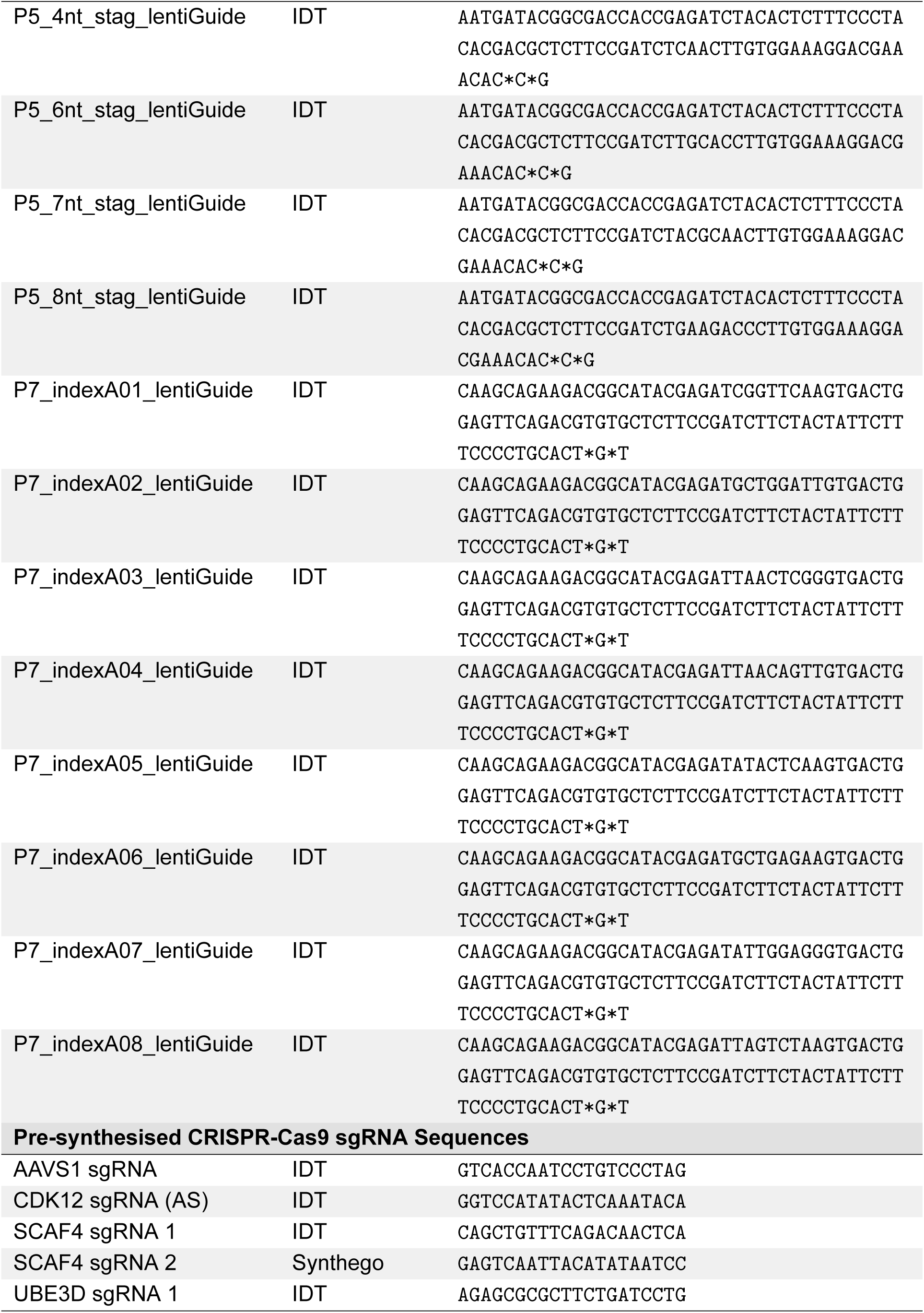
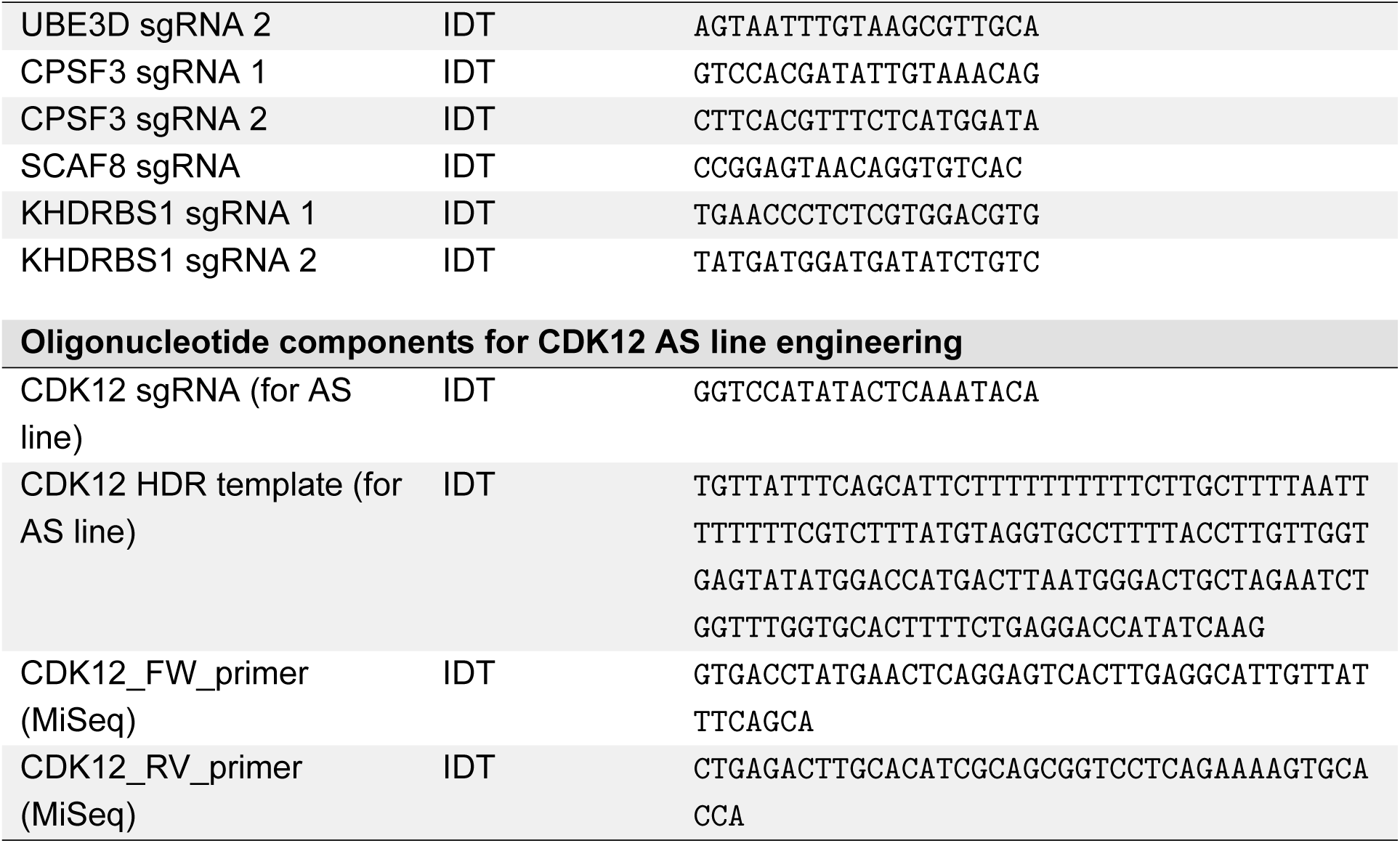
Oligonucleotides.

**Table 6.** Recombinant DNA.

| Plasmid Name | Source | Sequence (Link) |
| --- | --- | --- |
| CDK12_TurboID_handcloned.dna | cloned in-house | https://benchling.com/oliverozaydin/f/_nxnFF8WWt0-plasmids/ |
| SCAF4_complete_domainmutant.dna | cloned in-house | https://benchling.com/oliverozaydin/f/_nxnFF8WWt0-plasmids/ |
| SCAF4_onlyCID_domainmutant.dna | cloned in-house | https://benchling.com/oliverozaydin/f/_nxnFF8WWt0-plasmids/ |
| SCAF4_TurboID_handcloned.dna | cloned in-house | https://benchling.com/oliverozaydin/f/_nxnFF8WWt0-plasmids/ |
| SCAF4_ΔCID_domainmutants.dna | cloned in-house | https://benchling.com/oliverozaydin/f/_nxnFF8WWt0-plasmids/ |
| SCAF8_TurboID_handcloned.dna | cloned in-house | https://benchling.com/oliverozaydin/f/_nxnFF8WWt0-plasmids/ |

Table 6 continued from previous page
| Plasmid Name | Source | Sequence (Link) |
| --- | --- | --- |
| lentiGuide-puro.dna | Addgene: 52963 | https://benchling.com/oliverozaydin/f_nxnFF8WWt0-plasmids/ |
| pMSCV-IRES-mCherry.dna | Addgene: 52114 | https://benchling.com/oliverozaydin/f_nxnFF8WWt0-plasmids/ |
| pMSCV-IRES-eGFP.dna | Addgene: 20672 | https://benchling.com/oliverozaydin/f_nxnFF8WWt0-plasmids/ |
| PB_CAG_eGFP.dna | Addgene: 40973 | https://benchling.com/oliverozaydin/f_nxnFF8WWt0-plasmids/ |
| PB_CAG_tdTomato.dna | Addgene: 133569 | https://benchling.com/oliverozaydin/f_nxnFF8WWt0-plasmids/ |

**Table 7.** Software, command-line tools, and R packages used for data analysis.

| Tool/Package | Version | Purpose |
| --- | --- | --- |
| STAR | 2.7.11b | Alignment of RNA-seq reads to reference genomes |
| bowtie2 | 2.5.3 | Alignment of ChIP-seq reads to reference genomes |
| samtools | 1.19.2 | Processing, filtering, and indexing of BAM files |
| deeptools | 3.5.5 | Generation of bigwig files and signal matrices |
| MAGeCK | 0.5.9.5 | Statistical analysis of CRISPR screen data |
| TrimGalore | 0.6.10 | Quality and adapter trimming of raw sequencing reads |
| umi_tools | 1.1.6 | UMI extraction and deduplication of sequencing reads |
| FastQC / MultiQC | 0.12.1 / 1.24 | Quality control and report aggregation |
| Rsubread | 2.22.1 | Read alignment and feature counting |
| limma | 3.64.3 | Differential expression and linear modelling |
| edgeR | 4.6.3 | Differential expression analysis |
| APAlyzer | 1.22.0 | Alternative polyadenylation analysis |
| clusterProfiler | 4.16.0 | Functional enrichment analysis |
| GraphPad Prism | 11.0.0 | Statistical analysis of cell viability assays |
| FlowJo | 10.10.0 | Processing of FACS data |
| Affinity Designer | 2.6.2 | Figure collation and aesthetics |
| ggplot2 | 4.0.1 | Data visualisation |

Table 7 continued from previous page
| Tool/Package | Version | Purpose |
| --- | --- | --- |
| data.table | 1.18.0 | High-performance data manipulation |
| MASS / mgcv | 7.3.65 / 1.9.3 | Robust linear and generalised additive modelling |
| boot | 1.3.32 | Bootstrap confidence interval estimation |

**Table 8.** Cell lines and culturing conditions.

| Cell line | Cell type | Media composition | Culturing conditions & Source |
| --- | --- | --- | --- |
| HAP1 | *Homo sapiens* CML (derived from KBM-7) | IMDM medium, 10% FBS, 1X GlutaMAX (2mM), 100 U/mL Penicillin, 100 μM/mL Streptomycin | 37°C, 5% CO<sub>2</sub>. Horizon Discovery (C669) |
| MV-4;11 | *Homo sapiens* AML | RPMI 1640 medium, 10% FBS, 1X GlutaMAX (2mM), 100 U/mL Penicillin, 100 μM/mL Streptomycin | 37°C, 5% CO<sub>2</sub>. American Type Culture Collection (ATCC) (CRL-9591) |
| KBM-7 | *Homo sapiens* CML | IMDM medium, 10% FBS, 1X GlutaMAX (2mM), 100 U/mL Penicillin, 100 μM/mL Streptomycin | 37°C, 5% CO<sub>2</sub>. Horizon Discovery (C628) |
| HEK293T | *Homo sapiens* embryonic kidney | DMEM medium, 10% FBS, 1X GlutaMAX (2mM), 100 U/mL Penicillin, 100 μM/mL Streptomycin | 37°C, 5% CO<sub>2</sub>. American Type Culture Collection (ATCC) (CRL-3216) |
| A-375 | *Homo sapiens* melanoma | DMEM medium, 10% FBS, 1X GlutaMAX (2mM), 100 U/mL Penicillin, 100 μM/mL Streptomycin | 37°C, 5% CO<sub>2</sub>. American Type Culture Collection (ATCC) (CRL-1619) |
| NIH/3T3 | *Mus musculus* embryonic fibroblast | DMEM medium, 10% FBS, 1X GlutaMAX (2mM), 100 U/mL Penicillin, 100 μM/mL Streptomycin | 37°C, 5% CO<sub>2</sub>. American Type Culture Collection (ATCC) (CRL-1658) |
| Schneider's *Drosophila* line 2 (S2) | *Drosophila melanogaster* epithelial | Schneider's *Drosophila* Medium, 10% FBS, 1X GlutaMAX (2mM), 100 U/mL Penicillin, 100 μM/mL Streptomycin | Room temperature and atmospheric CO<sub>2</sub>. Thermo Fisher Scientific (R69007) |

### Method details

#### Analog-sensitive cell line generation

Analog-sensitive (AS) MV-4;11 CDK12, CDK12 and CDK12/CDK13 lines were provided as a gift from Prof. Ricky Johnstone (Peter MacCallum Cancer Centre). An additional CDK12-AS line was generated in-house in HAP1 cells by Cas9-mediated knock-in of F813G gatekeeper residue. Ribonucleoprotein (RNP) complexes were assembled by combining 116 pmol of CDK12-targeting sgRNA (IDT Alt-R CRISPR-Cas9 sgRNA or Synthego EasyEdit; 200 pmol/µL stock; sequence displayed in the Key Resources Table) with 39 pmol of Alt-R S.p. HiFi Cas9 Nuclease V3 (Integrated DNA Technologies, #1074182) and incubating at room temperature for 20 minutes prior to transfer to ice. 78 pmol of a single-stranded HDR donor encoding the F813G mutation (Integrated DNA Technologies, 100 pmol/µL stock; sequence displayed in the Key Resources Table) was added at this stage. 2 × 10^5^ HAP1 cells were resuspended in 20 µL of nucleofection solution (16.4 µL SE Nucleofector solution + 3.6 µL Supplement 1; SE Cell Line 4D-Nucleofector X kit, Lonza #V4XC-1032), combined with the assembled RNP in a 16-well Nucleocuvette strip kept on ice, and electroporated using the EN-138 program on a Lonza 4D-Nucleofector X Unit (Lonza, AAF-1003X). Electroporated cells were recovered overnight in pre-warmed IMDM in a 24-well plate prior to passaging into fresh medium the following day.

Single-cell clones were isolated by FACS sorting into 96-well plates containing IMDM supplemented with 20% (v/v) FBS using a BD FACSAria^TM^ Fusion (Becton Dickinson). Wells were inspected over 2–3 weeks for viability and confluency and validated for biallelic F813G knock-in by amplicon-based MiSeq sequencing of the CDK12 gatekeeper locus. Briefly, the gatekeeper region was amplified from extracted genomic DNA in a first-round PCR with locus-specific primers carrying universal Illumina 5′ overhangs (sequences displayed in the Key Resources Table), indexed in a second-round PCR with dual-barcoded primers, cleaned with AmPure XP beads (Beckman Coulter) at a 1:1 ratio, pooled at equimolar ratios, and sequenced on an Illumina MiSeq at 1 × 10^6^ reads per pool. This depth was sufficient to confidently call the binary outcome of biallelic knock-in.

#### Plasmid cloning

Any custom expression and tagging constructs used in this study were generated through Gibson assembly (constructs in the Plasmid Inventory table). Briefly, for domain mutant constructs, parental backbones were linearised by restriction digest using XbaI (NEB, #R0145S), BamHI-HF (NEB, #R3136S) or BsmBI-v2 (NEB, #R0739S) as appropriate (Chemicals, peptides, and recombinant proteins table), resolved on 0.8% (w/v) agarose gels run at 120 V for ≥40 minutes, gel-extracted using the Monarch Gel Purification Kit (NEB, #T1020S), and then quantified by NanoDrop (Thermo Scientific, #13-400-518). Gene inserts were synthesised by GenScript or Integrated DNA Technologies and amplified with NEBNext Ultra II Q5 Master Mix (NEB, #M0544S) using primer pairs (10 µM forward and reverse mix) carrying 15–30 nt overhangs complementary to the digested backbone, then gel-purified as above. Backbone and insert were combined with NEBuilder HiFi DNA Assembly Master Mix (NEB, #E5520S) in a 1:1 (volume) ratio and incubated at 50°C for 60 minutes, with insert-to-vector molar ratios set according to the manufacturer’s recommendations for the fragment sizes used. 1 *`µ`*L of the assembly reaction was used to transform chemically competent E. coli, plated onto LB-agar containing the appropriate antibiotic, and single colonies were expanded in liquid LB for plasmid extraction. Successful assemblies were verified by whole-plasmid Nanopore sequencing.

#### Lentiviral production

Lentiviral particles were produced in HEK293T cells using third-generation packaging plasmids pMDLg/pRRE (Addgene, #12251), pRSV-Rev (Addgene, #12253) and pMD2.G/VSV-G (Addgene, #12259) co-transfected with the relevant lentiviral transfer vector (Plasmid Inventory table). HEK293T cells were seeded in complete DMEM the day before transfection. Plasmids were co-transfected using Lipofectamine 3000 (Invitrogen, #L3000001) according to the manufacturer’s instructions, or with polyethyleneimine (PEI; Sigma-Aldrich, #03880-500ML) at a 3:1 (w/w) PEI:DNA ratio, and medium was refreshed 24 h post-transfection. At 48 h post-transfection, viral supernatant was harvested, passed through a 0.45 µm syringe filter to remove cellular debris, and concentrated by centrifugation through Amicon Ultra-15 centrifugal filter units (Millipore, #UFC903024). Concentrated virus was either snap-frozen in liquid nitrogen and stored at –80°C until use, or applied directly to target cells in the presence of 4 µg/mL Sequa-brene (Sigma-Aldrich, #S2667) to enhance transduction efficiency.

#### Cell viability assay

Cell viability was quantified using the CellTiter-Glo Luminescent Cell Viability Assay following small-molecule inhibitor treatment (Promega, #G7570). 5 × 10^3^ cells were seeded per well in 100 *`µ`*L of complete medium in opaque-walled 96-well plates (Corning Costar, #3917) to minimise luminescence bleed-through during readout. Inhibitors (3-MB-PP1 or SR-4835) were prepared at 20 *`µ`*M in complete medium and serially diluted 2-fold across the plate, with the final well reserved as a DMSO vehicle control; equal volumes of each inhibitor dilution and cell suspension were combined to yield a starting final concentration of 10 µM that decreased 2-fold in each successive well. Plates were incubated at 37°C for 72 h. On the day of readout, CellTiter-Glo reagent and substrate buffer (Promega) were equilibrated to room temperature for ≥30 min, plates were equilibrated to room temperature for a further 30 minutes, and CellTiter-Glo reagent was added at a 1:1 (v/v) ratio. Plates were mixed on an orbital shaker for 2 minutes to induce cell lysis, incubated for a further 10 minutes in the dark, and luminescence was recorded on a CLARIOstar Plus plate reader (BMG Labtech). A medium-only well receiving CellTiter-Glo reagent served as a background blank. All conditions were assessed in biological and technical triplicate, and dose-response curves were fitted with a four-parameter log-logistic model in GraphPad Prism (v11.0.0) to derive approximate IC_50_ values.

#### Genome-wide CRISPR–Cas9 screening

For genome-wide loss-of-function screening, HAP1, MV-4;11 and KBM7 cells stably expressing humanised S. pyogenes Cas9 were generated by lentiviral transduction with the FU-Cas9-Cherry vector (Addgene, #70182; gift from M. Herold), followed by FACS-based enrichment of the top quintile of mCherry+ cells on a BD FACSAria Fusion sorter. The Brunello sgRNA library (Addgene, #73178; gift from D. Root; Doench et al., 2016) was packaged in HEK293T cells across 20 × T150 flasks using PEI-based lentiviral production as described above, and viral supernatant was harvested, filtered, concentrated and aliquoted for screening. Optimal multiplicity of infection (MOI) was empirically determined for each cell line by transducing a dilution series of library virus and selecting with puromycin to identify the dose yielding 3̂0% cell survival relative to an untransduced, unselected control; an untransduced puromycin-treated negative control was included to confirm complete killing of non-integrated cells. Cells were transduced at the calibrated MOI (typically 0.3) to maintain a minimum library representation of ≥250 cells per sgRNA across all screens, with the cell number calculated as the product of the library size and the intended coverage divided by the intended MOI. Transduced cells were selected with 1 *`µ`*g/mL puromycin for 5–7 days, after which T0 baseline pellets were collected, and the remaining population was split into treatment arms (DMSO vehicle or small-molecule inhibitor at approximate IC_50_ derived from CellTiter-Glo assays). Cells were passaged for 14–21 days while maintaining ≥250-fold coverage at each passage, and T_end_ pellets were collected. All pellets were frozen on dry ice and stored at –80°C until processing.

Genomic DNA was extracted using the QIAamp DNA Blood Midi Kit (Qiagen, #51185) according to the manufacturer’s instructions. 7.5 *`µ`*g of gDNA was used per PCR reaction, with multiple reactions performed per sample and pooled post-amplification to ensure adequate representation of the integrated library. sgRNA cassettes were amplified using TaKaRa Ex Taq DNA Polymerase (Takara, #RR001A) in 50 *`µ`*L reactions with dNTPs, 1x ExTaq reaction buffer, 5 *`µ`*M staggered P5 forward primer mix and 5 *`µ`*M indexed P7 reverse primer (sequences in the Key Resources Table) under the following cycling conditions: 95°C for 1 min; 28 cycles of 95°C for 30 s, 53°C for 30 s and 72°C for 30 s; final extension at 72°C for 10 min. PCR products were cleaned using AmPure XP beads (Beckman Coulter, #A63881) at a 1:1 (v/v) bead-to-sample ratio with 5 min binding at room temperature, two washes in 70% (v/v) ethanol on a DynaMag magnetic rack and air-drying, before elution in 50 *`µ`*L of 1x TE buffer. Cleaned libraries were quantified, pooled at equimolar ratios and sequenced by the WEHI Genomics Core on an Illumina NextSeq 2000 (single-end, 100 cycles) at 100–400 × 10^6^ reads per run.

#### SDS–polyacrylamide gel electrophoresis and Western blotting

Whole-cell lysates were prepared by pelleting cells at 200 x g for 5 min, washing in ice-cold PBS, pelleting again, and resuspending the pellet directly in 2x Laemmli loading buffer (60 mM Tris-HCl pH 6.8, 10% (v/v) glycerol, 2% (w/v) SDS) at a 1:1 (v/v) ratio with residual PBS, followed by addition of *`β`*-mercaptoethanol and bromophenol blue from a 20x loading dye stock at 1:20 (v/v) final dilution. Lysates were denatured at 95°C for 10 min and either used immediately or stored at −80°C. Protein concentration was determined using the Qubit Protein Broad Range Assay Kit (Thermo Fisher Scientific, #Q33212) according to the manufacturer’s instructions, with diluted Laemmli buffer as the blank, and concentrations were multiplied by 1.05 to account for loading dye addition prior to loading. 20*`µ`*g of total protein was resolved on 4–15% Mini-PROTEAN TGX Precast Protein Gels (Bio-Rad, #4561086; 15-well, 15*`µ`*L) for routine analyses or on 7.5% Mini-PROTEAN TGX gels (Bio-Rad, #4561023; 10-well, 30 *`µ`*L) for proteins >150 kDa, in 1x Tris/Glycine/SDS running buffer (10% (v/v) dilution of 10x stock; Bio-Rad, #1610732) at 80 V for 20 min followed by 120 V for ≥1 h as required, alongside the Abcam Prestained Protein Ladder (Abcam, #ab116028; 10–245 kDa) or Precision Plus Protein Dual Color Standards (Bio-Rad, #1610374). Proteins were transferred to PVDF (Bio-Rad, #1704156; 0.2 *`µ`*m Mini) or nitrocellulose (Bio-Rad, #1704158; 0.2 *`µ`*m Mini) Trans-Blot Turbo transfer packs for chemiluminescent or fluorescent detection, respectively, using the Trans-Blot Turbo Transfer System (Bio-Rad, #1704150) on the mixed or high-molecular-weight pre-set programs as appropriate. Membranes were blocked in 5% (w/v) skim milk powder in TBS-T (Tris-buffered saline + 0.1% (v/v) Tween-20) for 1 h at room temperature on a tilting shaker, incubated with primary antibody diluted in 5% (w/v) skim milk/TBS-T overnight at 4°C, washed 4 x 5 min in TBS-T, incubated with the appropriate HRP-conjugated secondary antibody (in 5% (w/v) skim milk/TBS-T) for 1 h at room temperature, and washed 4 x 5 min again in TBS-T. Antibodies, suppliers and catalogue numbers are listed in the Antibodies table. Chemiluminescence was developed with SuperSignal West Pico PLUS Chemiluminescent Substrate (Thermo Fisher Scientific, #34580) and imaged on a ChemiDoc Imaging System (Bio-Rad, #12003153) at the highest resolution settings.

#### FACS-based competition assays

Cells used in competition assays were stably labelled with eGFP, mCherry, or tdTomato fluorescent reporters by PiggyBac transposition (PB-CAG-eGFP, Addgene #40973; PB-CAG-tdTomato, Addgene #133569) or lentiviral transduction (pMSCV-IRES-eGFP, Addgene #20672; pMSCV-IRES-mCherry, Addgene #52114). For each assay, competing cell populations were paired with opposing fluorescent markers (or one labelled and one unlabelled) and mixed 1:1 at assay commencement (T0). The founding fluorescent fraction was recorded by flow cytome-try on a NovoCyte Penteon (Agilent) before seeding. Mixed populations were cultured for 4-14 days with regular passaging in the presence of DMSO vehicle or small-molecule inhibitor at concentrations maintained constant across passages, and the eGFP: mCherry, eGFP: tdTomato, or labelled: unlabelled population ratio was recorded at each passage. Shifts in fluorescent proportions relative to T0 were used to quantify the relative fitness conferred by gene knockout, drug treatment, or their combination. All assays were performed in biological and technical triplicate at each time point and analysed in FlowJo (v10.10.0).

#### 3′ RNA sequencing library preparation

For 3’ mRNA sequencing experiments, adherent cell lines were seeded 16 h before treatment at 0.5 x 10^6^ cells/mL in 2 mL of complete medium per well of a 6-well plate, while suspension cell lines were seeded on the day of treatment at 1 x 10^6^ cells/mL under the same format. Following treatment, total RNA was harvested by direct addition of 500*`µ`*L of TRIzol Reagent (Thermo Fisher Scientific, #15596026) to the adherent monolayer (with the plate tilted during collection to minimise volume loss) or to suspension cell pellets recovered by centrifugation at 400 g for 5 min and removal of medium; lysates were homogenised by pipetting until cells had visibly dissociated and either processed immediately or stored at −80°C until extraction. Total RNA was purified using the Direct-zol RNA Miniprep Kit (Zymo Research, #R2052) according to the manufacturer’s instructions, with the addition of an equal volume of molecular-grade ethanol before loading on the column, centrifugation at 12000 g for 30 s, sequential washes with RNA Pre-Wash Buffer and RNA Wash Buffer, an additional dry spin to remove residual liquid, and elution with DNase/RNase-free water. RNA concentration was determined using either the Qubit RNA High Sensitivity Assay (Thermo Fisher Scientific, #Q33230) or the Qubit RNA Broad Range Assay (Thermo Fisher Scientific, #Q33265), depending on expected yield. Sequencing libraries were prepared from total RNA using the QuantSeq 3’ mRNA-Seq V2 Library Prep Kit with UMI Second Strand Synthesis Module (Lexogen, #191.96) according to the manufacturer’s instructions: an oligo(dT) primer carrying a linker sequence primes first-strand synthesis, the RNA template is degraded, second-strand synthesis incorporates a random primer carrying a unique molecular identifier (UMI), and the resulting double-stranded library is amplified with dual-indexed adapters by PCR before bead-based clean-up. Libraries were pooled at equimolar ratios and sequenced single-end on an Illumina NextSeq 2000. Each condition was sequenced in technical triplicate.

#### Transient transcriptome sequencing (TT-seq) library preparation

Cells were seeded the day prior to reach 70–80% confluency on the day of labelling (approximately 5.5 × 10^6^ HAP1 cells per 10 cm^2^ dish). Cells were labelled with 4-thiouridine (4sU; Sigma-Aldrich, #T4509) at a 1 mM final concentration (from a 1 M stock) for 15 min at 37°C, either alone (untreated control) or during the final 15 min of a 30 min exposure to medium containing the inhibitor of interest. Labelling was terminated by aspiration of medium and direct addition of 1 mL of TRIzol Reagent (Thermo Fisher Scientific, #15596026) while scraping the plate. After 4 min at room temperature, 200 *`µ`*L of chloroform was added with vigorous shaking for 15 s, samples were transferred to phase-separation tubes and centrifuged at 12,000×g for 15 min at 4°C, and the aqueous phase was precipitated with 1 volume of isopropanol (10 min, room temperature; 12,000×g, 20 min, 4°C), washed in 85% (v/v) ethanol, resuspended in DNase/RNase-free water, and denatured at 65°C for 10 min. RNA concentrations of experimental samples and of *Drosophila melanogaster* S2 4sU-labelled spike-in RNA were determined by Qubit RNA Broad Range Assay (Invitrogen, #Q10211), and each sample was adjusted to 5% (w/w) spike-in in a final volume of 100 *`µ`*L. 100 *`µ`*L samples were sonicated twice in microTUBE AFA Fiber screw-cap tubes (6 mm × 16 mm; Covaris, #520096) on a Covaris ME220 (30 s duration, 75 W peak power, 25% duty factor, 10,000 cycles per burst, 18.8 W average power, ∼6.4°C bath temperature), and fragmentation was confirmed by RNA ScreenTape on a TapeStation 4200 (Agilent, #G2991BA). 100 *`µ`*g of sonicated RNA was biotinylated in a 253 *`µ`*L reaction containing 100 *`µ`*L of DNase/RNase-free water, 3 *`µ`*L of biotin buffer (833 mM Tris-HCl pH 7.4, 83.3 mM EDTA), and 50 *`µ`*L of 0.1 mg/mL MTSEA biotin-XX (Biotium, #BT90066) reconstituted in dimethylformamide, at 25°C for 30 min on a Thermomixer in the dark. Unreacted biotin was removed by two sequential phenol:chloroform:isoamyl alcohol (25:24:1; Thermo Fisher Scientific, #15593031) extractions through Phase-Lock-Gel tubes (12,000×g, 5 min, 4°C). Biotinylated RNA was precipitated with 0.1 volumes of 5 M NaCl and 1.1 volumes of isopropanol (10 min, room temperature; 20,000×g, 20 min, 4°C), washed in 85% (v/v) ethanol (20,000×g, 5 min, 4°C), and resuspended in 52 *`µ`*L of DNase/RNase-free water. Biotinylation efficiency was confirmed by slot blot: 1:20 dilutions of each sample were applied to a Hybond-N membrane (Cytiva, #RPN203N) on a slot blot apparatus fitted with three sheets of filter paper (Cytiva, #3030-917) under vacuum, UV-crosslinked twice at 0.2 J/cm^2^ (254 nm) on a Stratalinker 2400, blocked in 10% (w/v) SDS, 1 mM EDTA in PBS for 20 min with agitation, probed with HRP-conjugated streptavidin (Thermo Fisher Scientific, #N100; 1:50,000) for 15 min at room temperature, washed sequentially in 1% and 0.1% (w/v) SDS in PBS (10 min each), developed with SuperSignal West Pico PLUS Substrate (Thermo Fisher Scientific, #34580), and counter-stained with methylene blue (0.5 M sodium acetate pH 5.2, 0.5% (w/v) methylene blue) for 5 min followed by overnight destaining in water to verify uniform RNA loading.

For enrichment, RNA was denatured at 65°C for 10 min, cooled on ice for 5 min, and incubated with 200 *`µ`*L of *`µ`*MACS Streptavidin MicroBeads (Miltenyi, #130-074-101) per 100 *`µ`*g of labelled RNA on a rotator for 30 min at room temperature. The mixture was loaded onto *`µ`*Columns pre-rinsed with 100 *`µ`*L of nucleic acid equilibration buffer on a *`µ`*MACS separator, washed twice with 500 *`µ`*L of wash buffer (100 mM Tris-HCl pH 7.4, 10 mM EDTA, 1 M NaCl, 0.1% (v/v) Tween-20) pre-warmed to 65°C, and labelled RNA was eluted by two sequential additions of 100 *`µ`*L of freshly prepared 100 mM DTT in RNase-free water (5 min apart). Eluates were cleaned using the RNeasy MinElute Cleanup Kit (Qiagen, #74204) according to the manufacturer’s instructions, except that 1050 *`µ`*L of molecular-grade ethanol was added per reaction to retain a broad range of fragment sizes, and final RNA was eluted in 17 *`µ`*L of DNase/RNase-free water with concentration assessed by Qubit RNA High Sensitivity Assay (Invitrogen, #Q32852). Sequencing libraries were prepared using the NEBNext Ultra II Directional RNA Library Prep Kit for Illumina (NEB, #E7760L) following the rRNA-depleted FFPE RNA protocol of the manufacturer’s instructions, with library integrity and concentration assessed by TapeStation 4200 RNA ScreenTape and Qubit RNA High Sensitivity Assay prior to pooling and sequencing.

#### DRB-TT-seq library preparation

DRB\TT-seq was performed to assess the elongation rate of RNAPII in the same manner as the TT-seq workflow with the following modifications. Cells were pre-incubated with 100 *`µ`*M DRB (Sigma-Aldrich, #D1916) for 3.5 hours to block transcription elongation before release. To remove residual DRB and initiate synchronous transcription elongation wave release, cells were quickly washed twice with pre-warmed PBS before the addition of pre-warmed media containing the respective treatments. To capture early temporal elongation wave processivity kinetics, samples were treated and harvested at 0, 15, and 30 minutes post-release. For all conditions, 4SU was added to the media at a final concentration of 1 mM during the final 15 minutes of the treatment window before harvest. Following collection, RNA extraction, fragmentation, biotinylation, enrichment, and strand-specific library preparation were executed identically to the standard TT-seq workflow described above.

#### Immunoprecipitation–mass spectrometry (IP-MS)

For IP-MS experiments targeting SCAF4, KHDRBS1 or RNAPII, HAP1 cells were seeded in quadruplicate 15 cm dishes and grown to 70–90% confluency, then treated with 10 *`µ`*M SR-4835 or an equivalent volume of DMSO for 90 min at 37°C prior to collection. Cells were washed twice with ice-cold PBS and subjected to multi-step subcellular fractionation as previously described^6^, with all buffers supplemented with PhosSTOP (Merck, #4906845001) and cOmplete EDTA-free Protease Inhibitor Cocktail (Merck, #04693132001). Cell pellets were resuspended in ≥10x pellet volumes of hypotonic wash buffer (10 mM HEPES pH 7.5, 10 mM KCl, 1.5 mM MgCl_2_) and incubated on ice for 45 min. Nuclei were pelleted at 1000 g for 15 min at 4°C, washed once in hypotonic buffer and re-pelleted at 1000 g for 15 min at 4°C. Nuclei were resuspended in 2x pellet volumes of nucleoplasmic lysis buffer (20 mM HEPES pH 7.9, 1.5 mM MgCl_2_, 150 mM potassium acetate, 10% (v/v) glycerol, 0.05% (v/v) NP-40) and incubated on ice for 20 min, after which chromatin was pelleted at 20000 g for 20 min at 4°C and the supernatant was collected as the nucleoplasmic fraction. The chromatin pellet was resuspended in 2x pellet volumes of low-salt chromatin digestion buffer (20 mM HEPES pH 7.9, 1.5 mM MgCl_2_, 10% (v/v) glycerol, 0.05% (v/v) NP-40, 150 mM NaCl, benzonase at 1:1000) and incubated on ice for 60 min; following centrifugation at 20000 g for 20 min at 4°C, the low-salt supernatant was pooled with the nucleoplasmic fraction. The remaining undigested chromatin was extracted in 2x pellet volumes of high-salt chromatin digestion buffer (20 mM HEPES pH 7.9, 3 mM EDTA, 1.5 mM MgCl_2_, 10% (v/v) glycerol, 500 mM NaCl, 0.1% (v/v) NP-40) on ice for 20 min, diluted with 6x pellet volumes of high-salt dilution buffer (20 mM HEPES pH 7.9, 3 mM EDTA, 1.5 mM MgCl_2_, 10% (v/v) glycerol, 0.1% (v/v) NP-40), centrifuged at 20000 g for 20 min at 4°C, and the supernatant was pooled with the prior nuclear fractions. Pooled nuclear lysates were then used for immunoprecipitation.

Lysates were quantified by BCA assay (Pierce) and normalised to approximately 500 *`µ`*g of protein per IP. 2 *`µ`*g of either rabbit or mouse IgG control, anti-RNAPII (CTD4H8; Merck, #05-623), anti-SCAF4 (Bethyl, #A303-951A) or anti-SCAF8 (Bethyl, #A301-036A) (Antibodies table) was added per sample and incubated overnight at 4°C with end-over-end rotation. Protein A/G beads were washed 2x in PBS, resuspended in IP wash buffer (150 mM NaCl, 20 mM Tris-HCl pH 7.5, 1.5 mM MgCl_2_, 3 mM EDTA, 10% (v/v) glycerol, 0.1% (v/v) NP-40), added to each sample and incubated at 4°C with end-over-end rotation for 4 h. Beads were washed 5x in IP wash buffer, and immunoprecipitated proteins were eluted by incubation in 50 *`µ`*L of elution buffer (100 mM Tris-HCl pH 7.5, 1% (w/v) SDS, 0.5 mM EDTA) at 65°C for 3 min, followed by two further 30 *`µ`*L elutions performed identically. Pooled eluates were snap-frozen prior to S-Trap clean-up. Protein digestion was carried out using S-Trap Micro spin columns (ProtiFi, #K002-MICRO-0010KT) according to the manufacturer’s protocol: SDS was added to each eluate to a final concentration of ≥2% (w/v), disulfide bonds were reduced and alkylated by simultaneous addition of TCEP (5 mM final) and chloroacetamide (CAA; 20 mM final) at 55°C for 30 min, and samples were acidified with phosphoric acid to ∼2.5% final concentration. Acidified lysates were diluted 1:10 in S-Trap binding buffer (100 mM TEAB pH 7.5 in 90% (v/v) methanol) and loaded onto S-Trap Micro columns by centrifugation at 4000 g for 1 min, with the flow-through discarded. Columns were washed three times in binding buffer with 180° rotation between washes, and proteins were digested on-column with 1 *`µ`*g of sequencing-grade trypsin in 50 mM ammonium bicarbonate at 47°C for 2 h with the column loosely capped. Peptides were sequentially eluted at 4000 g for 1 min in 50 mM TEAB, then 0.2% (v/v) formic acid, and finally 50% (v/v) acetonitrile, and pooled eluates were dried by vacuum centrifugation prior to LC-MS/MS.

Peptides were separated by reverse-phase liquid chromatography on a 15 cm C18 fused-silica column with an integrated emitter tip (IonOpticks; 75 *`µ`*m ID, 360 *`µ`*m OD, 1.6 *`µ`*m C18 beads) on a Thermo Scientific NeoVanquish LC system coupled to a Thermo Scientific Orbitrap Eclipse Tribrid mass spectrometer, using a 34 min linear gradient from 2% to 34% buffer B (80% (v/v) acetonitrile, 0.1% (v/v) formic acid). Data were acquired in data-independent acquisition (DIA) mode with MS1 settings of 124000 resolution, m/z 375–1500 scan range and 300% AGC target, and tMSn settings of HCD collision energies of 24, 28 and 32%, m/z 200–1200 scan range, automatic maximum injection time and custom AGC target. Raw DIA data were searched in Spectronaut (v19.9)^59^ using directDIA mode with BGS factory settings, carbamidomethylation of cysteine as a fixed modification, and N-terminal acetylation and methionine oxidation as variable modifications. The result filter m/z range was set to 300–1800 with a relative intensity threshold of 5.

#### Proximity labelling TurboID

For TurboID proximity-labelling experiments, HAP1 cells stably expressing CDK12-TurboID, SCAF4-TurboID, or SCAF8-TurboID were seeded in 15 cm dishes in DMEM supplemented with dialysed FBS and 1x GlutaMAX, and grown to 70–90% confluency. Cells were treated for 60 min at 37°C with 10 *`µ`*M SR-4835 or an equivalent volume of DMSO, after which biotin was added to a final concentration of 100 *`µ`*M and labelling was allowed to proceed at 37°C for 30 min. Cells were then washed 5x with ice-cold PBS to quench biotinylation, collected by manual scraping, and subjected to multi-step subcellular fractionation as described above for the IP-MS workflow.

Lysates were quantified by BCA assay (Pierce) and protein quantities were normalised across samples before incubation with high-capacity streptavidin agarose beads overnight at 4°C with end-over-end rotation. Beads were washed 3x in 0.5% (w/v) SDS in PBS, proteins were reduced by incubation in 100 mM DTT in 0.5% (w/v) SDS in PBS, and beads were washed 2x in UC buffer (6 M urea, 100 mM Tris-HCl pH 8.5). Beads were transferred to Pierce Snap-Cap Spin Columns (Thermo Fisher Scientific, #69725) and alkylated for 20 min in 50 mM iodoacetamide in UC buffer in the dark. Beads were then washed by 1 min spins at 1000 g sequentially with UC buffer (6x), PBS (4x) and MS-grade ultrapure water (3x). Beads were resuspended in 50 mM ammonium bicarbonate (NH_4_HCO_3_) containing 2 *`µ`*g of sequencing-grade trypsin and digested overnight at 37°C. Digested peptides were collected by centrifugation at 1000 g for 1 min, beads were washed once with 50 mM ammonium bicarbonate, and the eluates were pooled, dried at 45°C in a SpeedVac, and resolubilised in mass spectrometry loading buffer (2% (v/v) acetonitrile, 0.1% (v/v) formic acid) prior to LC-MS/MS.

Mass spectrometry data acquisition (LC parameters, 34 min linear gradient, DIA acquisition on a Thermo Scientific Orbitrap Eclipse Tribrid coupled to a NeoVanquish LC system) and Spectronaut directDIA search were performed identically to the IP-MS workflow described above.

#### Co-immunoprecipitation

For Co-IP experiments, HAP1 CDK12^AS/AS^ cells were seeded in quadruplicate 15 cm dishes in IMDM supplemented with 10% (v/v) FBS and 1x GlutaMAX, and grown to 70–90% confluency. Cells were washed 2x with ice-cold PBS and subjected to multi-step subcellular fractionation as described above for the IP-MS workflow^6^, with the following modifications: cell pellets were resuspended in 10x pellet volumes of hypotonic wash buffer (10 mM HEPES pH 7.5, 10 mM KCl, 1.5 mM MgCl_2_) and incubated on ice for 25 min, and the supernatant recovered after centrifugation at 1000 g for 15 min at 4°C was retained as the cytoplasmic fraction rather than discarded. The remaining nuclear pellet was sequentially extracted into nucleoplasmic, low-salt chromatin digestion and high-salt chromatin digestion buffers as described for IP-MS, with the high-salt extract diluted in 6x pellet volumes of salt-free dilution buffer (20 mM HEPES pH 7.9, 3 mM EDTA, 1.5 mM MgCl_2_, 10% (v/v) glycerol, 0.1% (v/v) NP-40) prior to clarification at 20000 g for 20 min at 4°C, and the three resulting supernatants were pooled to form the final nuclear fraction.

After fractionation, both cytoplasmic and nuclear lysates were quantified by BCA assay (Pierce) and normalised to approximately 800 *`µ`*g of protein per IP. Protein A magnetic beads (Pierce) were washed 2x in PBS and resuspended in IP wash buffer (150 mM NaCl, 20 mM Tris-HCl pH 7.5, 1.5 mM MgCl_2_, 3 mM EDTA, 10% (v/v) glycerol, 0.1% (v/v) NP-40); 25 *`µ`*L of beads and 3.5 *`µ`*g of either rabbit IgG (Bethyl Laboratories), rabbit anti-SCAF4 (Bethyl, #A303-951A) or rabbit anti-CPSF3 (ABclonal, #A2222) were added per sample and incubated with end-over-end rotation at 4°C overnight. Beads were washed 5x in IP wash buffer, and immunoprecipitated proteins were eluted by addition of SDS loading dye and boiling at 95°C for 10 min prior to SDS-PAGE and Western blot analysis.

#### Chromatin immunoprecipitation sequencing (ChIP-seq) library preparation

For ChIP-seq, Protein A/G magnetic beads (Thermo Fisher Scientific, #88848) were pre-blocked by 2x washes in ChIP buffer (20 mM Tris-HCl pH 8.0, 150 mM NaCl, 2 mM EDTA, 1% (v/v) Triton X-100, 0.15% (w/v) SDS) followed by incubation in blocking buffer (ChIP buffer plus 0.1% (w/v) BSA) at 4°C with end-over-end rotation until use, to minimise non-specific chromatin binding. HAP1 cells were seeded the day prior at ∼1 x 10^7^ cells per 15 cm dish to reach ∼2 x 10^7^ cells on the day of crosslinking. Cells were crosslinked in freshly prepared crosslinking buffer (50 mM HEPES-KOH pH 7.5, 100 mM NaCl, 1 mM EDTA, 0.5 mM EGTA, 11% (v/v) formaldehyde) diluted 1:10 into cell medium to give a final formaldehyde concentration of 1% (v/v), for 10 min at room temperature with rocking; the reaction was quenched by addition of 125 mM glycine, and cells were pelleted at 500 g for 5 min at 4°C.

Nuclei were enriched in nuclear enrichment buffer (20 mM Tris-HCl pH 8.0, 10 mM NaCl, 2 mM EDTA, 0.5% (v/v) NP-40 substitute) supplemented with PhosSTOP (Merck, #4906845001) and cOmplete EDTA-free Protease Inhibitor Cocktail (Merck, #04693132001) at 1 tablet per 50 mL, and resuspended in 1 mL of sonication buffer (20 mM Tris-HCl pH 7.5, 150 mM NaCl, 2 mM EDTA, 1% (v/v) NP-40 substitute, 0.3% (w/v) SDS). Chromatin was fragmented on a Covaris ME220 Focused Ultrasonicator (Covaris, #500506) for 8 min per sample at 75 W peak power, 15% duty factor, 1000 cycles per burst (11.3 W average power), with NIH/3T3 cells processed in parallel under identical conditions to provide a cross-species spike-in. Solubilised chromatin was clarified by maximum-speed centrifugation at 4°C for 20 min, diluted 1:1 in ChIP dilution buffer (20 mM Tris-HCl pH 8.0, 150 mM NaCl, 2 mM EDTA, 1% (v/v) Triton X-100) and quantified, with NIH/3T3 chromatin added at exactly 5% of total DNA content per sample and 1% of each sample reserved as input. Each IP combined 30 *`µ`*g of fragmented chromatin with 10 *`µ`*g of primary antibody (anti-RNAPII clone 4H8, Merck #05-623; or anti-RNAPII pSer2 clone 3E10, Active Motif #61083), 50 *`µ`*L of pre-blocked Protein A/G beads (diluted 1:1 in ChIP buffer) and 20 *`µ`*L of 10% (w/v) BSA (0.1% (w/v) final), in a final volume of 2 mL adjusted with ChIP dilution buffer, and incubated overnight at 4°C with end-over-end rotation.

Beads were collected on a magnet and washed sequentially at 4°C with low-salt ChIP buffer (20 mM Tris-HCl pH 8.0, 150 mM NaCl, 2 mM EDTA, 1% (v/v) Triton X-100, 0.15% (w/v) SDS), high-salt wash buffer (20 mM Tris-HCl pH 8.0, 500 mM NaCl, 2 mM EDTA, 1% (v/v) Triton X-100, 0.1% (w/v) SDS), LiCl wash buffer (20 mM Tris-HCl pH 8.0, 250 mM LiCl, 2 mM EDTA, 0.5% (v/v) NP-40, 0.5% (w/v) sodium deoxycholate) and TE buffer (10 mM Tris-HCl pH 7.5, 1 mM EDTA), with 5 min of end-over-end rotation per wash. DNA was eluted and proteins were digested by incubation in 150 *`µ`*L of reverse-crosslinking buffer containing 300 *`µ`*g/mL proteinase K at 55°C for 1 h; supernatants were transferred and decrosslinked overnight at 65°C alongside input controls. DNA was purified using the ChIP DNA Clean & Concentrator Kit (Zymo Research, #D5205) according to the manufacturer’s instructions, with two sequential elutions in 11 *`µ`*L of pre-warmed (55°C) elution buffer. DNA concentration and fragment size were assessed using the Qubit dsDNA High Sensitivity Assay (Invitrogen, #Q32854) and TapeStation 4200 D1000 ScreenTape (Agilent, #G2991BA), respectively. Sequencing libraries were prepared using the NEBNext Ultra II DNA Library Prep Kit for Illumina (NEB, #E7645L) according to the manufacturer’s instructions, assessed by TapeStation for fragment size distribution and yield, pooled at equimolar ratios, and sequenced paired-end on an Illumina NextSeq 2000.

#### Time lapse imaging

Cell confluence was assayed every 1 hour by time-lapse imaging using the IncuCyte Live Cell Analysis imaging (Essen BioScience) for 72 hours with 5% CO2 and 37°C climate control. Briefly, 4x 10^5^ CDK12AS HAP1 cells were nucleofected with 5uM negative control antisense oligonucleotide (NC-ASO) or 5*`µ`*M U1 antisense oligonucleotide (U1-ASO). 24 hours post nucleofection, 1x 10^4^ cells were seeded in a 96-well plate and treated with DMSO or 3MBPP1 (5*`µ`*M). The plate was then transferred to the IncuCyte system and imaged for 72 h. Cell confluence was calculated based on the imaged area using the in-built ”Basic Analyzer” modulepost-nucleofection.

#### Phosphoproteomic mass-spectrometry

Enrichment of phosphorylated peptides was performed based on the EasyPhos protocol^60^. HAP1 CDK12^AS/AS^ cells were treated with 10 *`µ`*M 3-MB-PP1 or an equivalent volume of DMSO for 20 min at 37°C, washed on ice with 100 mM Tris-HCl pH 8.5, and lysed by addition of chilled SDC lysis buffer (4% (w/v) sodium deoxycholate, 100 mM Tris-HCl pH 8.5). Cells were scraped, heat-treated at 95°C for 5 min, sonicated with a tip probe, and clarified. Protein quantities were determined by BCA assay (1:50 CuSO_4_:bicinchoninic acid) and normalised to 350 *`µ`*g per sample. Proteins were reduced with 10 mM TCEP and alkylated with 40 mM 2-chloroacetamide (CAA) at 45°C for 5 min with shaking (1500 rpm), then digested overnight to peptides with trypsin and LysC (1:160 enzyme:protein w/w) in 0.1% (v/v) acetic acid at 37°C with shaking (1500 rpm).

For phosphopeptide enrichment, isopropanol (50% (v/v) final), trifluoroacetic acid (TFA; 6% (v/v) final) and KH_2_PO_4_ (1 mM final) were added to the digests, vortexed, and clarified at 16000 g for 5 min. Supernatants were transferred to a deep-well plate containing 12 mg of TiO_2_ beads in 100% acetonitrile (ACN) and shaken at 45°C and 1500 rpm for 5 min. Beads were sedimented at 2000 g for 1 min, the supernatant was aspirated, and beads were washed five times in 5% (v/v) TFA / 60% (v/v) isopropanol before being transferred onto in-house C8 Stage Tips in 0.1% (v/v) TFA / 60% (v/v) isopropanol and spun to dryness at 1500 g. Phosphopeptides were eluted from the beads through the stage tip with 32% (v/v) ACN / 5.6% (w/v) ammonium hydroxide (NH_4_OH) at 1500 g, dried in a SpeedVac at 45°C, and resuspended in 0.1% (v/v) TFA / 99% (v/v) isopropanol. The eluate was loaded onto an in-house SDB-RPS Stage Tip, spun to dryness at 1500 g, washed sequentially with 0.1% (v/v) TFA / 99% (v/v) isopropanol and 0.2% (v/v) TFA / 5% (v/v) ACN, and eluted with 0.14% (w/v) NH_4_OH / 60% (v/v) ACN. Eluates were dried at 45°C in a SpeedVac and resuspended in mass spectrometry loading buffer (2% (v/v) ACN, 0.1% (v/v) formic acid).

Peptides were separated by reverse-phase liquid chromatography on a 15 cm C18 fused-silica column with an integrated emitter tip (IonOpticks; 75 *`µ`*m ID, 360 *`µ`*m OD, 1.6 *`µ`*m C18 beads) on a Thermo Scientific Ultimate 3000 RSLC Nano-LC system coupled via an Easy-nLC source to a Thermo Scientific Orbitrap Eclipse Tribrid mass spectrometer (Thermo Fisher Scientific, #FSN04-10000). Peptides were loaded at 600 nL/min in buffer A (99.9% (v/v) Milli-Q water, 0.1% (v/v) formic acid) and eluted at 400 nL/min over a 30 min linear gradient from 2% to 34% buffer B (90% (v/v) acetonitrile, 0.1% (v/v) formic acid). Data were acquired in data-independent acquisition (DIA) mode: MS1 spectra were collected in the Orbitrap at 120000 resolution with a normalised AGC target of 300%, custom maximum injection time, RF lens at 40%, and a scan range of m/z 375–1500 in profile mode, with dynamic exclusion of 30 s applied to all charge states of each precursor; MS2 spectra were collected in the Orbitrap at 30000 resolution following HCD fragmentation with normalised collision energy, scan range m/z 200–1200, custom AGC target and automatic maximum injection time, in profile mode.

### Quantification and statistical analysis

#### CRISPR screen analysis

Demultiplexed FASTQ files were trimmed of adaptor and stagger sequences using cutadapt (v4.9) and quality-checked with FastQC (v0.12.1) and MultiQC (v1.24). sgRNA reads were counted against the Brunello library reference and tested for enrichment using MAGeCK (v0.5.9.5), with the rra subcommand for pairwise time-resolved comparisons and the mle subcommand for multi-condition analyses. T0 baseline samples were required to have a Gini index < 0.1 to confirm uniform library representation prior to selection. Gene-level β-scores (MLE) and log fold-changes with RRA scores (RRA) were used to rank candidate hits, and visualisation was performed in R (v4.5.2) using ggplot2 (v4.0.1).

#### 3’ RNA sequencing analysis

UMIs were extracted from raw reads using umi_tools extract (v1.1.6) and reads were aligned to GRCh38/hg38 using the Rsubread align function (v2.22.1) with default single-end parameters. Aligned BAM files were deduplicated with umi_tools dedup to remove PCR duplicates while preserving unique transcript counts, and strand-specific bigWig tracks were generated using deeptools bamCoverage (v3.5.5) with --bin-size 10 --normalizeUsing CPM. Gene-level read counts were quantified using the featureCounts function (Rsubread v2.22.1) with annot.inbuilt = “hg38” and strandSpecific = 1^61^. Count matrices were imported into edgeR (v4.6.3), filtered for lowly expressed genes using filterByExpr and normalised by trimmed mean of M-values (TMM) using calcNormFactors^62^ . Differential expression was tested using the limma-voom pipeline (voom, lmFit, contrasts.fit, eBayes; limma v3.64.3) with experimental contrasts defined per comparison^63^.

#### Intronic polyadenylation (IPA) analysis with APAlyzer

For intronic polyadenylation (IPA) quantification, UMI-extracted reads were aligned to GRCh38/ hg38 using STAR (v2.7.11b) with --sjdbGTFfile set to the GENCODE v49 primary assembly annotation, deduplicated with umi_tools dedup, and strand-specific bigWig tracks were generated using deeptools bamCoverage (v3.5.5) with --bin-size 10 --normalizeUsing CPM. IPA events were quantified using the APAlyzer package (v1.22.0)^41^ by mapping reads against a custom hg38 polyadenylation site (PAS) reference (hg38_REF.RData; https://github.com/R JWANGbioinfo/PAS_reference_RData) using the REF4PAS function, with intronic read densities quantified using the PASEXP_IPA function. The relative expression (RE) of each intronic PAS was calculated as the log_2_-ratio of the difference between intronic upstream and intronic downstream read densities relative to the constitutive 3’ terminal exon read density, with a pseudocount of *`ɛ`* = 0.1 added to all densities before log transformation to minimise the effect of low-count regions. Differential IPA usage was tested in edgeR (v4.6.3) on raw intronic upstream counts after filterByExpr and TMM normalisation^62^, using a Quasi-Likelihood negative binomial generalised linear model fitted with glmQLFit and evaluated with glmQLFTest on experimental contrasts. Difference-of-differences contrasts were used to capture the rescue effect of target knockout on CDK12 inhibition-induced IPA, yielding a relative expression difference (RED) per IPA event calculated as the difference between knockout and AAVS1 control conditions in their CDK12i-induced RE changes; positive RED values indicate enhanced IPA usage in the knockout upon CDK12 inhibition, while negative values indicate rescue.

#### Transient transcriptome (TT) sequencing analysis

Raw FASTQ files were quality- and adaptor-trimmed using trim_galore (--quality 20 --length 20) and aligned to a hybrid human (GRCh38/hg38) and *Drosophila melanogaster* (dm6) reference genome using STAR (v2.7.11b) with --twopassMode Basic --outFilterMultimapNmax 20 --outSAMattributes All. PCR duplicates were marked using picard-tools (v3.3.0) Mark- Duplicates, and BAM files were filtered with samtools (v1.19.2) to retain high-quality alignments (MAPQ ≥ 30), excluding unmapped, secondary, and duplicate reads (-F 1284). For stranded dUTP libraries, reads were separated by strand using --filterRNAstrand. Spike-in normalisation factors were computed as the ratio of a 10^6^ scaling constant to the number of uniquely aligned dm6 reads, and applied to correct for global differences in nascent transcription between samples.

Signal matrices for metagene and transcription dynamics analyses were generated using computeMatrix (deeptools v3.5.5) at three scales: all protein-coding genes scaled to a 15 kb gene body with 5 kb flanking regions; long genes (> 100 kb) scaled to a 100 kb gene body to assess transcriptional processivity; and windows centred on intronic polyadenylation (IPA) sites identified by 3’ RNA-seq (±4 kb, --binSize 20 --referencePoint center). Metagene profiles were computed separately for sense and antisense strands, stratified by gene length quartile, and summarised using 5% trimmed means with bootstrap confidence intervals (n = 200 iterations); profiles were smoothed with a Savitzky-Golay filter (signal package, p = 3, n = 31).

To quantify transcription termination and processivity defects at the 3’ ends of long genes (> 100 kb), a terminal signal metric was computed from deepTools-derived sense-strand matrices. The terminal signal was defined as the log_2_-transformed mean signal across the final 25% of the scaled gene body (bins 326–400) with a pseudocount of 0.1 added to the term; genes with a mean terminal signal ≤ 0.1 were excluded to minimise stochastic effects of noise. CDK12 inhibition-sensitive target genes displaying severe processivity defects were dynamically isolated by filtering for genes that exhibited a terminal signal decrease of log_2_ fold-change < –0.5 upon CDK12 inhibition (3MB treatment) relative to the vehicle control (DMSO) in wild-type cells. To evaluate the rescue of these termination defects by genetic or pharmacological perturbation, the treatment-induced change in terminal signal (ΔTerminal Signal = 3MB – DMSO) was calculated across this defective gene cohort for both control and perturbed backgrounds. Statistical differences in the magnitude of transcription termination failure and subsequent rescue between groups were evaluated using two-sided, paired Wilcoxon signed-rank tests.

#### DRB/TT-sequencing analysis

DRB/TT-seq data were processed as described for TT-seq with the following modifications. Duplicate reads were retained rather than removed, as biological duplicates may arise from synchronised RNAPII in this assay context; all other quality filters were applied identically (MAPQ ≥ 30, excluding unmapped and secondary reads). Bigwig files were generated using CPM normalisation (deeptools bamCoverage, --normalizeUsing CPM --binSize 10) rather than spike-in normalisation, as the DRB release assay is used to measure wavefront progression rather than steady-state transcriptional output. Signal matrices were computed from the TSS to 150 kb downstream using computeMatrix (--afterRegionStartLength 100000 --binSize 100 --referencePoint TSS). Genes were required to exceed 100 kb in length and possess detectable signal in the wild-type DMSO 30-minute release condition (mean per-bin signal ≥ 0.1 in the first 500 bins) to be considered candidates for wavefront analysis.

Per-bin signal (S_i_) was z-scored relative to an unlabelled control distribution (mean *µ*, standard deviation *σ*) with a pseudocount of 0.001 to prevent division by zero and suppress stochastic noise:

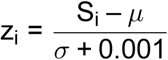

The transcriptional wavefront was identified by applying a rolling mean smoother (k = 50 bins) to the z-score signal vector; the wavefront position was defined as the first bin at which the smoothed z-score fell below 0.5. Genes for which no signal depletion was detected were assigned the maximum window value. Wavefront positions were additionally required to advance monotonically between timepoints. Elongation rate was calculated as the displacement of the wavefront from the TSS (d_t_) over the release duration (t):

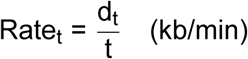

Metagene profiles were displayed as z-scores relative to the untreated control group rather than spike-in normalised signal, with 95% confidence intervals derived by bootstrapping (n = 200 iterations) and smoothing applied using a Savitzky-Golay filter (p = 3, n = 21).

ChIP-sequencing analysis

Raw FASTQ files were quality- and adaptor-trimmed using trim_galore (v0.6.10) with --quality 20 --illumina --length 20 --paired and aligned to a hybrid human (GRCh38/hg38) and mouse (mm10) reference genome using bowtie2 (v2.5.3) with parameters --very-sensitive, --no-discordant, and --no-mixed. BAM files were filtered with samtools (v1.19.2) to retain high-quality paired alignments (MAPQ ≥ 30), with PCR duplicates and non-primary alignments removed (-F 1024 and -F 264, respectively). Spike-in normalisation factors were calculated using DESeq2 based on the ratio of total mapped hg38 to mm10 reads to correct for global differences in RNAPII occupancy across conditions, and applied during bigwig generation via the --scaleFactor parameter of deeptools bamCoverage (v3.5.5); additional parameters included --binSize 10 --smoothLength 30 --extendReads.

Signal matrices for metagene analysis were generated using computeMatrix scale-regions (deeptools v3.5.5), scaling protein-coding genes to a 15 kb gene body with 5 kb flanking regions up- and downstream of the TSS and TES (--skipZeros --missingDataAsZero --binSize 50). Metagene profiles were restricted to long protein-coding genes (> 100 kb), summarised using 5% trimmed means with 95% confidence intervals derived by bootstrapping (n = 200 iterations), and smoothed with a third-order Savitzky-Golay filter (sgolayfilt).

#### Immunoprecipitation mass spectrometry and proximity labelling TurboID analysis

Raw MS data were searched with Spectronaut v19.9. Precursor-level data from the second-pass search were used for downstream analysis, with precursors exhibiting a Global Q-value or Library Q-value exceeding 0.01 excluded. Non-proteotypic precursors, single-precursor protein groups, and compound protein groups were additionally removed during pre-processing, and retained precursor intensities were log_2_-transformed. Differential expression analysis was performed using the dpcDE pipeline in limpa (v1.1.6)^64^, following protein-level summarisation with dpcQuant, which models missing values using detection probability curves (DPC)^64,65^. A 5% FDR threshold was applied to define significant hits. TurboID mass spectrometry data were acquired and analysed using identical parameters.

#### Phosphoproteomic mass-spectrometry analysis

Raw DIA data were searched in Spectronaut (v19.9)^59^ using directDIA mode with BGS factory settings, with serine/threonine/tyrosine phosphorylation, N-terminal acetylation and methionine oxidation as variable modifications and cysteine carbamidomethylation as a fixed modification. PTM localisation filtering was enabled with a minimum localisation threshold of 0. Result filter m/z was set to 300-1800 with a relative intensity threshold of 5, and DIA identification thresholds (precursor Q-value cutoff, precursor PEP cutoff, protein Q-value cutoff at experiment and run level, and protein PEP cutoff) were all set to 0.01 against the *Homo sapiens* reference proteome. PTM sites were quantified using the dpcQuantByRow function in limpa (v1.1.7), followed by differential analysis using the dpcDE function. A false discovery rate (FDR) threshold of 5% was used to define differentially phosphorylated sites.

## Supporting information

Supplementary Table 1 (Genome-wide CRISPR screen data)

## ACKNOWLEDGEMENTS

This research was supported by the Snow Medical Research Foundation [SMRF2021-SF346]. S.J.V was supported by a CSL Centenary Fellowship and a Snow Medical Fellowship; O.O was supported by an Australian Government Research Training Scholarship. J.R.D. and R.W.J. were supported by the National Health and Medical Research Council (NHMRC) (Ideas Grant and Investigator Grant, respectively), Worldwide Cancer Research, the Leukaemia & Lymphoma Society (R.W.J.), The Kids’ Cancer Project (R.W.J.), and the Barrie Dalgleish Centre for Myeloma and Associated Blood Diseases (R.W.J.). M.L. and G.K.S were supported by an NHMRC Investigator Grant (2025645). T.H.B was supported by Australian Research Council (ARC) Centre of Excellence Grant. The funders were not involved in the design of the study, collection, analysis, and interpretation of the data, the writing of this report, or the decision to submit the article for publication. We greatly thank the Walter and Eliza Hall Institute (WEHI) Genomics, Proteomics and FACS Facilities for sample and data collection (Vineet Vaibhav and Laura Dagley for the proteomics facility). We thank all members of the Vervoort lab for critical discussions and technical support throughout the project.

## Author contributions

S.J.V. conceived and designed the study. O.O., L.L., D.G., C.G., S.R., L.C., B.P., Y.T., and T.N. developed and established the experimental methodology. L.L., O.O., D.G., C.G., S.R., L.C., M.H., B.P., M.S., Y.T., and T.N. performed the investigations and conducted the core experiments. O.O., S.R., M.L. and B.P. processed the raw genomic sequencing data and executed the bioinformatics pipelines. O.O. performed the formal statistical and data analysis. O.O. and L.L. curated and maintained the data resources. Z.F., J.R.D., R.W.J., C.K., G.S., J.J., and T.B. provided reagents and experimental advice. O.O. and S.J.V wrote the original draft. O.O., S.J.V, and L.L. reviewed and edited the manuscript. S.J.V supervised the study and provided scientific oversight. S.J.V served as the project administrator and acquired the funding.

## Data, code and materials availability

- This study did not generate any novel reagents or code, but desired material or reagents may be requested through the corresponding author.
- All genomics and proteomics data will be made accessible in the Gene Expression Omnibus (GEO) and PRoteomics IDEntifications (PRIDE) databases; respectively. Uncompressed raw imaging data will be made accessible via Mendeley Data.
- Any additional requests for resources, reagents or information required to reanalyse the data reported in this paper should be directed to the corresponding author, Dr. Stephin Vervoort.

## SUPPLEMENTARY FIGURES

**Supplementary Figure 1.**
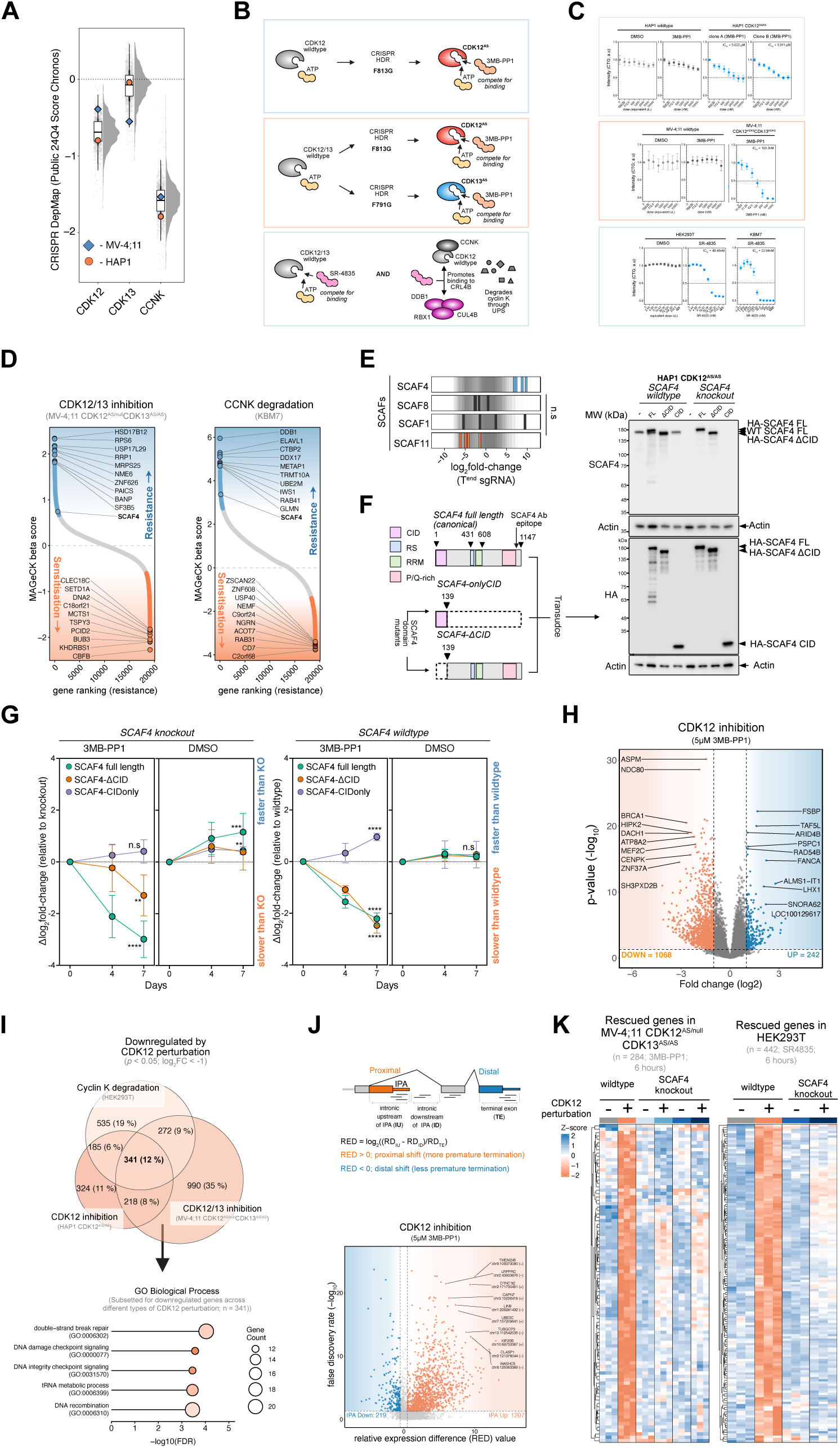
(A) Cancer Dependency Map (Public 24Q4 Score; Chronos) scores across cell-types for CDK12, CDK13 and CCNK. Orange circles represent HAP1 cells, while blue diamonds represent MV-4;11 cell. HAP1 cells were selected as the primary CDK12 analog-sensitive model for genome-wide screening applications owing to the measurable CDK12 dependency and experimental tractability of this context. (B) Schematic displaying mode of action for CDK12 analog-sensitive cells treated with 3MB-PP1 (top), CDK12/13 analog-sensitive treated with 3MB-PP1 (bottom) and SR-4835 treated cells. (C) Cell viability readings (Cell-Titre Glo®) following 3MB-PP1 treatment in HAP1 CDK12^AS/AS^ cells (top), 3MB-PP1 treatment in MV-4;11 CDK12^AS/null^ CDK13^AS/AS^ (middle) and SR-4835 treatment in HEK293T or KBM7 cells (bottom) across the dose curve following 72 hours of treatment. IC_50_, values in the top left with dots corresponding to the mean and error bars (*±*SEM). (D) Genes ranked from highest to lowest MAGeCK *β* scores from genome-wide knockout screens in MV-4;11 CDK12^AS/null^ CDK13^AS/AS^ (left) and KBM7 cells (right) following CDK12 perturbation. The colour of points is indicative of significant resistance (blue) and sensitisation hits (orange). (E) Rugplot displaying log_2_fold-change in sgRNA abundance across SR-related CTD associated factors following CDK12 inactivation in HAP1 CDK12^AS/AS^ cells. (F) Left: Schematic of domain-mutants leveraged for competitive proliferation assays indicating SCAF4 domains with absent regions encased in dotted boxes. Right: Western blot validation of domain-mutant expression in cell lines leveraged for competitive proliferation assays with *β*-actin as a loading control. (G) Competitive proliferation assay for HAP1 CDK12^AS/AS^ knockout (left) and wildtype (right) cells expressing domain mutants as indicated. Significance values were calculated through one-way ANOVA and are represented as follows: ns: p > 0.05, *: p *≤* 0.05, **: p *≤* 0.01, ***: p *≤* 0.001, ****: p *≤* 0.0001. Performed in biological and technical triplicate. (H) Volcano plot displaying differentially expressed genes in HAP1 CDK12^AS/AS^ cells treated with 5μM 3MB-PP1 for 6 hours. Significantly up-regulated (*n = 242*) and down-regulated (*n = 1068*) genes are based on a log_2_FC > |1| and adjusted p value (p_adj_ < 0.05). (I) Top: Venn diagram displaying intersection between downregulated genes across different cell-types and forms of CDK12 perturbations (p < 0.05, log_2_FC < -1). Bottom: Over-representation analysis of conserved downregulated genes across CDK12 perturbations (n = 341). (J) Top: 3’ RNA-seq data was analyzed with APAlyzer which calculates a value of relative expression difference (RED) by comparing read densities upstream of known IPA sites relative to reads from terminal exons^41^. Bottom: Volcano plot displaying false-discovery rate (-log_10_) versus relative expression difference (RED) in HAP1 CDK12^AS/AS^ cells treated with 5*µ*M 3MB-PP1. (K) Rescued genes following SCAF4 knockout in MV-4;11 CDK12^AS/null^ CDK13^AS/AS^ (left) and HEK293T (right) cells are displayed in subsequent heatmaps with cell colour corresponding to z-score log_2_FC per replicate. Performed in technical triplicate.

**Supplementary Figure 2.**
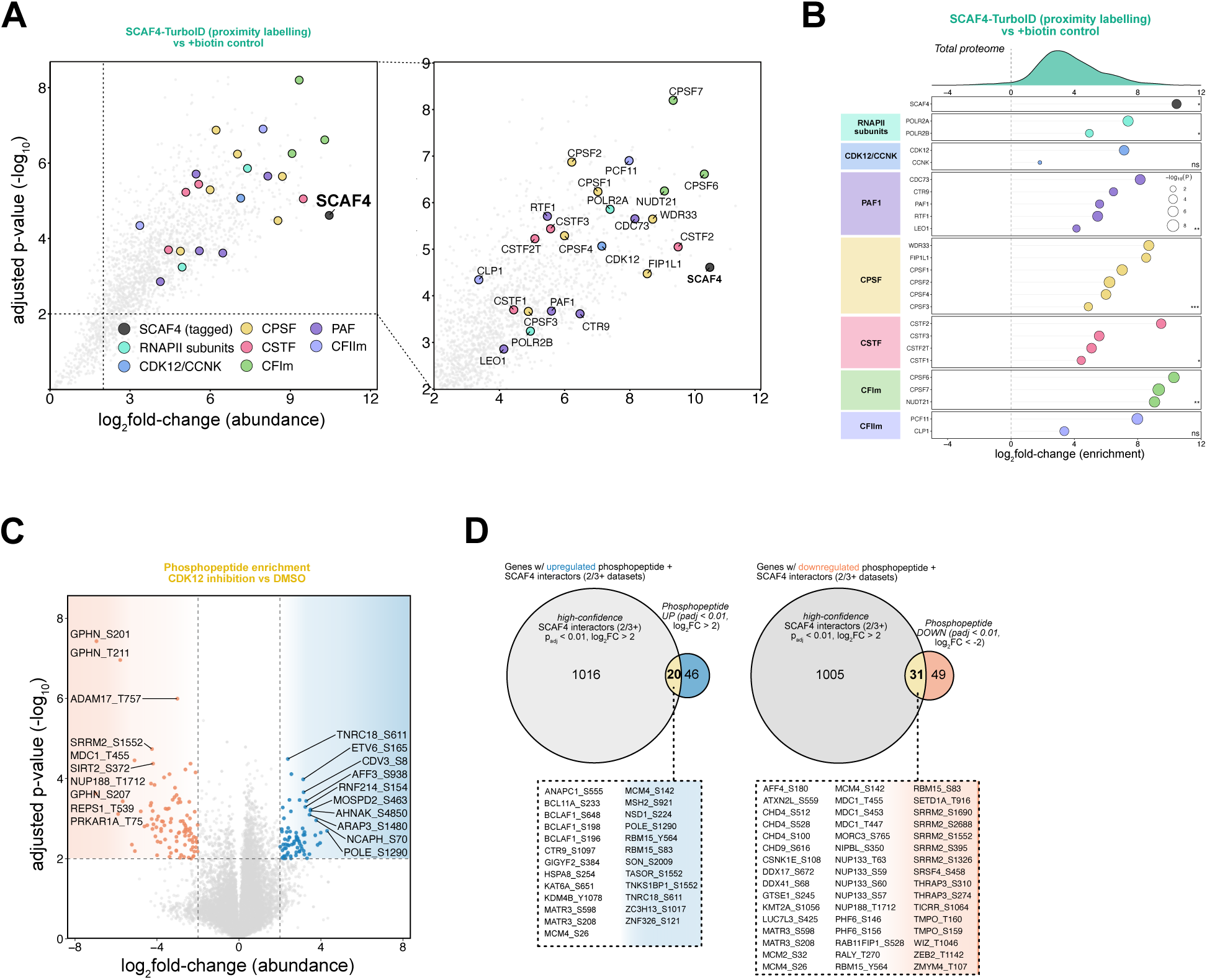
(A) Volcano plots mapping SCAF4 proximity-interactome identified via SCAF4-TurboID compared to biotin control. Complexes are highlighted by colour with zoomed-in region displaying interaction partner names. (B) Lollipop plot grouped by complex member enrichment in SCAF4 Turbo-ID data. log_2_fold-change in abundance is indicated on the x-axis, and circle size is indicative of statistical significance. (C) Volcano plot displaying differentially enriched phosphopeptides in HAP1 CDK12^AS/AS^ cells treated with 5μM 3MB-PP1 for 20 min. Significantly up-regulated and down-regulated sites are based on a (log_2_FC > |2| and p_adj_ < 0.01). (D) Venn diagram displaying intersection between upregulated (left) and downregulated (right) phosphosites and high-confidence SCAF4 interactors derived from IP-MS and TurboID assays (log_2_FC > |2| and p_adj_ < 0.01).

**Supplementary Figure 3.**
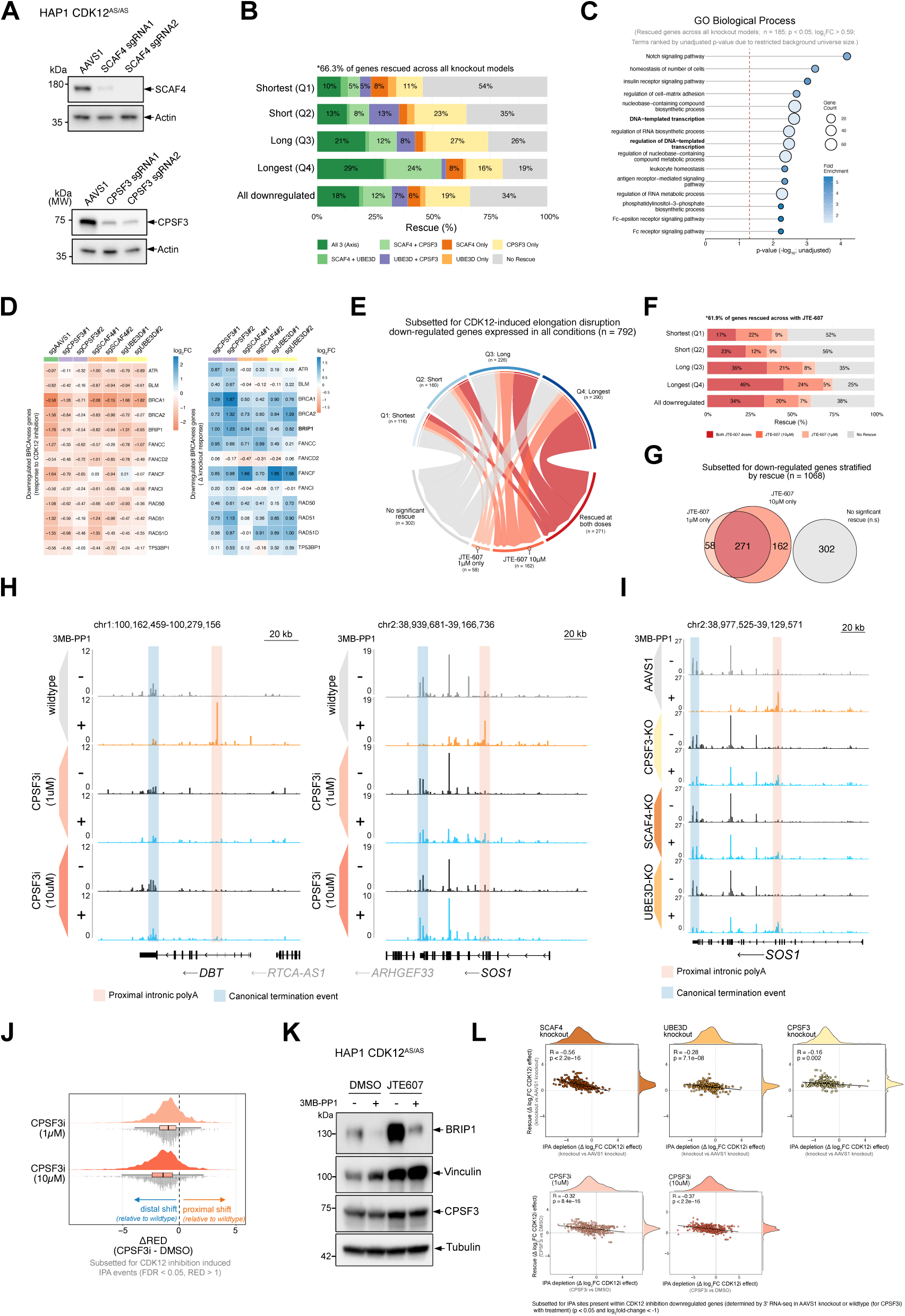
(A) Western blot validation of SCAF4 and CPSF3 knockout efficiency in HAP1 CDK12^AS/AS^ cells using two independent sgRNAs per target with actin serving as a loading control. (B) Stacked bar plot displaying the percentage rescue of downregulated genes within shortest to longest gene length quartiles (Q1–Q4) following knockout of the indicated factors. (C) Over-representation gene ontology (GO) biological process enrichment analysis of genes rescued across all knockout models (n = 185) ranked by unadjusted p-value, with circle size indicative of gene count and colour aligned with fold enrichment. (D) Heatmaps displaying log_2_fold changes amongst a curated set of CDK12 inactivation sensitive homologous recombination and DNA repair genes (BRCAness) genes across different knockout backgrounds (left) and Δlog2FC values following knockout (right) (E) Chord diagram of down-regulated genes upon CDK12 inhibition (n = 1,010), grouped by gene length quartile and rescue status following acute treatment with CPSF3 inhibitor JTE-607 at 1*µ*M or 10*µ*M. (F) Stacked bar plot displaying the percentage rescue of downregulated genes across gene length quartiles (Q1–Q4) following chemical inhibition with JTE-607 at 1*µ*M or 10*µ*M. Performed in technical triplicate. (G) Venn diagram showing the overlap of down-regulated genes rescued by 1*µ*M or 10*µ*M JTE-607 treatment. (H) Genome browser tracks showing normalized 3*^′^* RNA-seq signal intensity across the *DBT* (left) and *SOS1* (right) gene loci under DMSO and CPSF3 inhibitor (CPSF3i) treatment conditions *±* 3MB-PP1 5*µ*M . (I) Genome browser tracks showing normalized 3*^′^* RNA-seq signal intensity across *SOS1* (right) loci following different knockout conditions *±* CDK12 inhibition. (J) Raincloud plots showing ΔRED (Relative Expression Difference) values for CDK12 inhibition-induced IPA events (FDR < 0.05, RED > 1) following JTE-607 treatment relative to vehicle control (DMSO). ΔRED reflects changes in polyadenylation site usage, with negative and positive values indicating distal and proximal shifts, respectively. (K) Western blot showing BRIP1 expression levels following DMSO or 5*µ*M JTE-607 treatment *±* CDK12 inhibition; Vinculin, CPSF3, and Tubulin serve as loading controls. (L) Scatter plots displaying the correlation between IPA changes (ΔRED) and gene-level rescue (Δlog_2_ fold changes) across SCAF4, UBE3D, and CPSF3 knockout models as well as CPSF3i treatments.

**Supplementary Figure 4.**
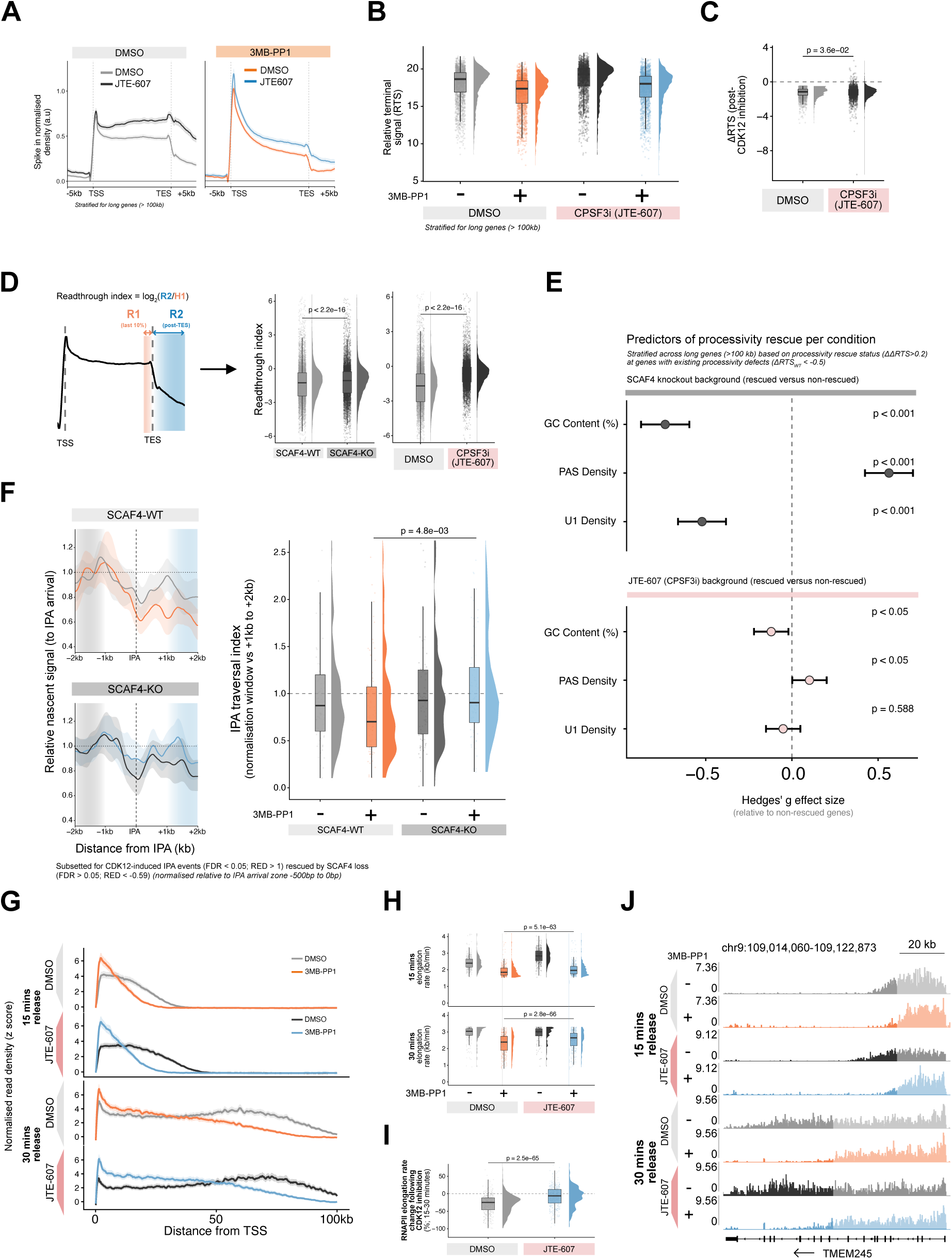
(A) Metagene profiles showing spike-in normalised TT-seq signal intensity across long gene bodies (> 100 kb) in DMSO or JTE-607 (10μM) treated cells treated with either DMSO or 3MB-PP1 (10μM). (B) Raincloud plots showing relative termination signal values across long genes (> 100 kb) in DMSO and JTE-607 treated cells *±* CDK12 inhibition (right). (C) Raincloud plots displaying change in RTS (ΔRTS) to evaluate transcription processivity following CDK12 inhibition in wildtype and SCAF4 knockout backgrounds. p-value was calculated through a two-sample, paired Wilcoxon signed-rank test. (D) Left: Schematic displaying readthrough index calculation; Right: Readthrough index value per condition were compared through a two-sample, paired Wilcoxon signed-rank test. (E) Hedges’ g effect size plots identifying gene-level architectural features (GC content, PAS density, and U1 density) and their contribution to processivity rescue status (ΔΔRTS > 0.2) in SCAF4-KO and JTE-607-treated backgrounds. (F) Left: Metagene profiling of relative nascent signal at CDK12-inhibition induced IPA events (FDR < 0.05, RED > 1 that were rescued by SCAF4 loss in IPA analysis. Right: IPA traversal index was calculated by comparing the nascent RNA signal within the immediate IPA arrival zone (-500bp to 0bp) to a downstream window (1000bp to 2000bp). Statistical testing between groups was performed using a Wilcoxon rank-sum test. (G) Metagene profiles displaying normalised nascent RNA read density (z-score) as a function of distance from the TSS after 15 or 30 minutes of DRB-release in indicated conditions. (H) Raincloud plots indicating RNAPII elongation rates (kb/min) at 15 min (top) and 30 min (bottom) post-DRB release under DMSO (–) or CDK12-inhibited (+) conditions stratified by treatment with JTE-607 or vehicle control (DMSO). p-values calculated using a two-sample, paired Wilcoxon signed-rank test. (I) Raincloud plots representing the percentage change in RNAPII elongation rates between the 15- and 30-minute intervals following CDK12 inhibition stratified by treatment with JTE-607 or vehicle control (DMSO). p-value calculated using a two-sample, paired Wilcoxon signed-rank test.. (J) Representative genome browser tracks at the *TMEM245* locus demonstrating nascent RNA wavefront progression stratified by timepoint and treatment with JTE-607 or vehicle control (DMSO) *±* CDK12 inhibition.

**Supplementary Figure 5.**
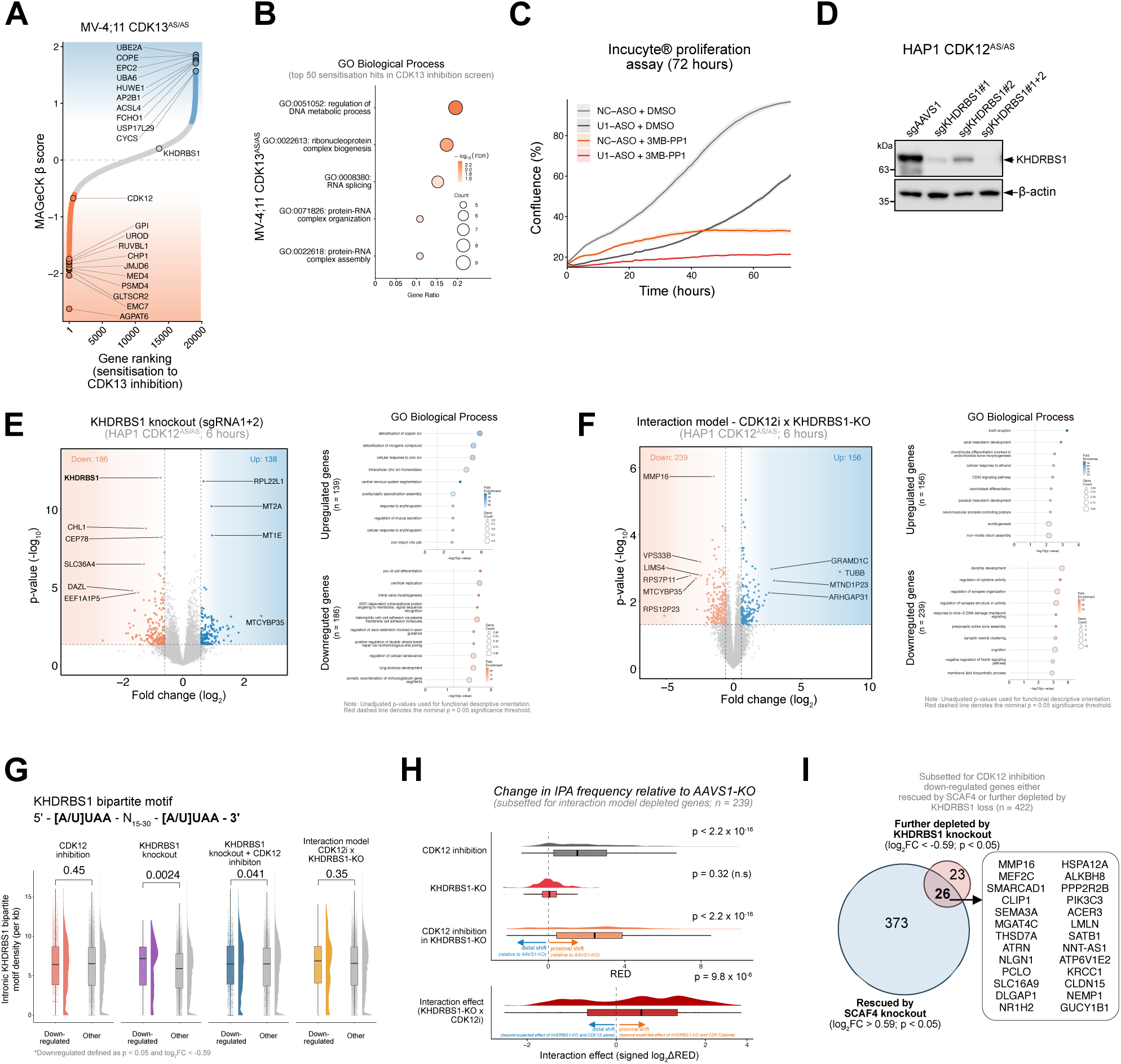
(A) Gene ranking from lowest to highest MAGeCK *β* score from a genome-wide CRISPR knockout screen in MV-4;11 CDK13^AS/AS^ cells following CDK13 inhibition with 7*µ*M 3MB-PP1 over 21 days. (B) Over-representation analysis of top 50 sensitisation candidates from the MV-4;11 CDK13^AS/AS^ screen where circle size represent gene count and circle colour represents FDR (-log_10_). (C) Incucyte proliferation assay measuring HAP1 cell confluence over 72 hours following treatment with 5*µ*M U1 snRNP antisense oligonucleotides (U1-ASO) or non-targeting control (NC-ASO) under DMSO or 3MB-PP1 (5*µ*M) conditions. (D) Western blot validation of KHDRBS1 knockout efficiency in HAP1 cells using two independent single-guide RNAs (sgRNA1 and sgRNA2) and an AAVS1 control guide (sgAAVS1); *β*-actin served as the loading control. (E) Volcano plot displaying differentially expressed genes (log_2_FC > |0.59| and p < 0.05) in HAP1 CDK12^AS/AS^ KHDRBS1-KO cells following 6 hours of CDK12 inhibition with 5*µ*M of 3MB-PP1 (left), coupled with GO enrichment profiles for the up- and down-regulated gene cohorts (right). (F) Volcano plot of interaction-model statistics highlighting genes showing specific synergistic disruption (orange) or rescue (blue) following combined KHDRBS1 loss and CDK12 inactivation (log_2_FC > |0.59| and p < 0.05) in HAP1 CDK12^AS/AS^ KHDRBS1-KO cells (left), coupled with GO enrichment profiles for the rescued (upregulated) and further-depleted (downregulated) gene cohorts (right). (G) Box plots indicating intronic KHDRBS1 bipartite motif density (5*^′^*-[A/U]UAA-N_15-30_-[A/U]UAA-3*^′^*) across downregulated versus non-downregulated gene cohorts across the indicated experimental conditions. (H) Ridge plots showing the net change in intronic polyadenylation frequency (ΔRED) relative to AAVS1-KO controls under single (top) or synergistic effects of CDK12 inhibition and KHDRBS1 knockout at IPAs within further depleted genes (n = 239). (I) Venn diagram highlighting the overlap of CDK12 inactivation downregulated that are either rescued by SCAF4 loss, further depleted by KHDRBS1 or co-regulated by both.

